# The adaptation dynamics between copy-number and point mutations

**DOI:** 10.1101/2022.08.11.503575

**Authors:** Isabella Tomanek, Calin C. Guet

## Abstract

Copy-number and point mutations form the basis for most evolutionary novelty through the process of gene duplication and divergence. While a plethora of genomic sequence data reveals the long-term fate of diverging coding sequences and their *cis*-regulatory elements, little is known about the early dynamics around the duplication event itself. In microorganisms, selection for increased gene expression often drives the expansion of gene copy-number mutations, which serves as a crude adaptation, prior to divergence through refining point mutations. Using a simple synthetic genetic system that allows us to distinguish copy-number and point mutations, we study their early and transient adaptive dynamics in real-time in *Escherichia coli*. We find two qualitatively different routes of adaptation depending on the level of functional improvement selected for: In conditions of high gene expression demand, the two types of mutations occur as a combination. Under low gene expression demand, negative epistasis between the two types of mutations renders them mutually exclusive. Thus, owing to their higher frequency, adaptation is dominated by copy-number mutations. Ultimately, due to high rates of reversal and pleiotropic cost, copy-number mutations may not only serve as a crude and transient adaptation, but also constrain sequence divergence over evolutionary time scales.

## Introduction

Adaptive evolution proceeds by selection acting on mutations, which are often implicitly equated with point mutations, that is, changes to a single nucleotide in the DNA sequence. However, nature is full of different types of bigger-scale mutations, such as mutations to the copy number of genomic regions ranging from only a few base pairs up to half a bacterial chromosome (Anderson and Roth, 1977; Darmon and Leach, 2014). The specific properties of mutations, such as their rate of formation and reversal, might influence the evolutionary dynamics in major ways, but are rarely considered. In bacteria, which are our focus, the duplication of genes or genomic regions occurs orders of magnitude more frequently than point mutations, ranging from 10^−6^ up to 10^−2^ per cell per generation (Roth *et al*., 1988; Drake *et al*., 1998; Andersson and Hughes, 2009; Elez *et al*., 2010; Reams and Roth, 2015). Moreover, while duplications can form via different mechanisms, they all are genetically unstable (Andersson and Hughes, 2009); the repeated stretch of DNA sequence is prone to *recA*-dependent homologous recombination. At rates between 10^−3^ and 10^−1^ per cell per generation duplications will reverse to the single copy (deletion) or duplicate further (amplification) (Roth *et al*., 1988; Andersson and Hughes, 2009; Mats E. Pettersson *et al*., 2009; Reams and Roth, 2015; Tomanek *et al*., 2020). Amplification of a gene or genomic region will, to a first approximation, increase its expression by means of elevated gene dosage (Elde *et al*., 2012; Gruber *et al*., 2012; Näsvall *et al*., 2012; Yona, Frumkin and Pilpel, 2015; Steinrueck and Guet, 2017; Belikova *et al*., 2020; Todd and Selmecki, 2020). Not surprisingly, due to their high rate of formation, gene amplifications are adaptive in situations where a rapid increase in gene expression is needed: resistance to antibiotics, pesticides or drugs via over-expression of resistance determinants (Prody *et al*., 1989; Albertson, 2006; Bass and Field, 2011; Nicoloff *et al*., 2019), immune evasion (Belikova *et al*., 2020) or novel metabolic capabilities through increased expression of spurious enzymatic side-activities (Blount *et al*., 2020; Richts *et al*., 2021). Due to their high intrinsic rate of deletion, often combined with significant fitness cost (Bergthorsson, Andersson and Roth, 2007; Mats E Pettersson *et al*., 2009; Reams *et al*., 2010), copy-number mutations not only differ from point mutations in their frequency of occurrence, but also in the nature of their reversibility.

Together, copy-number and point mutations are responsible for the evolution of most functional novelty of genes through the process of duplication and divergence of existing genes (Ohno, 1970; Kacser and Beeby, 1984; Conant and Wolfe, 2008; Andersson *et al*., 2015). Owing to the dynamic nature of gene duplication formation and reversal, the interplay between copy-number and point mutations may lead to complex evolutionary dynamics around the time point of origin of a new gene duplication. However, so far most attention has been focused on understanding the long-lasting process of how duplicate gene pairs diverge by accumulating point mutations (Lynch and Conery, 2000; Teufel, Masel and Liberles, 2015; Friedlander *et al*., 2017), while we know little about the potentially short-lived initial duplication event itself (Innan and Kondrashov, 2010). On one hand, this bias is due to significant technical challenges in studying transient copy-number variation experimentally (Andersson and Hughes, 2009; Lauer and Gresham, 2019; Belikova *et al*., 2020; Tomanek *et al*., 2020), and on the other hand, research has focused on the plethora of long-term evolutionary data that document the sequence divergence of paralogs, as “attention is shifted to where the data are” (Kondrashov, 2012).

In bacteria adaptive amplification, that is, amplification as a response to selection as opposed to neutral duplication and divergence, is considered the default mode of paralog evolution (Andersson and Hughes, 2009; Treangen and Rocha, 2011; Copley, 2020) and has been conceptualized in the Innovation-Amplification-Divergence (IAD) model (Bergthorsson, Andersson and Roth, 2007), which was later validated by evolution experiments (Elde *et al*., 2012; Näsvall *et al*., 2012). The IAD model posits that selection for a novel enzymatic activity leads to adaptive gene amplification that increases expression of an existing enzyme if it exhibits low levels of a beneficial secondary enzymatic activity (also referred to as promiscuous functions (Aharoni *et al*., 2005; Tawfik, 2010; Copley, 2017)). Eventually, protein sequences diverge as point mutations improve the secondary enzymatic function: a new protein function is born from an existing one. After the new (improved) function is present, superfluous additional gene copies will be lost due to their cost and high rate of reversibility, leaving only the copies of the two (ancestral and evolved) paralogs (Bergthorsson, Andersson and Roth, 2007; Reams *et al*., 2010; Elde *et al*., 2012; Näsvall *et al*., 2012). Similarly, adaptive amplification can precede the divergence of promoter sequences under selection favoring increased gene expression (Steinrueck and Guet, 2017). Thus, gene amplifications serve as a fast adaptation which can later be replaced by point mutations either within the coding region of a gene, increasing a cryptic enzymatic activity, or in its non-coding promoter region, increasing its expression (Elde *et al*., 2012; Näsvall *et al*., 2012; Yona, Frumkin and Pilpel, 2015; Steinrueck and Guet, 2017).

Since elevated numbers of gene copies provide an increased target for point mutations to occur (San Millan *et al*., 2017) it has been suggested that copy-number mutations speed up the process of divergence (Andersson and Hughes, 2009). However, if both, copy-number and point mutations are adaptive (Gruber *et al*., 2012) they also have the potential to interact epistatically. This interaction could result in unexpected evolutionary dynamics due to the different rates of formation and reversal of the two different mutation types.

To fill the knowledge gap that exists at around ‘time zero’ of the duplication-divergence process (Innan and Kondrashov, 2010) we designed a synthetic genetic system with which we can monitor, in real time, arising copy-number and point mutations in evolving populations of *Escherichia coli*. Importantly, while our results are also relevant to the divergence of paralogous protein sequences, here we study the process of divergence in a model gene promoter. Our genetic reporter system allows us to phenotypically distinguish between copy-number and point mutations, by specifically selecting for the increased expression of an existing but barely expressed gene. With our system at hand, we set out to test whether adaptive copy-number mutations facilitate or hinder adaptation by point mutation.

## Results

The motivation for this work was sparked by an evolution experiment conducted in *E*.*coli* at a locus exhibiting high rates of gene amplification (Steinrueck and Guet, 2017), which failed to produce any evolved clones with point mutations and thus lead us to hypothesize that copy-number mutations may interfere with the evolution by point mutations under certain conditions.

### An experimental system that distinguishes copy-number and point mutations

To study the interplay between point and copy-number mutations during adaption, we follow the fate of a barely expressed gene during its evolution towards higher expression. Our experimental system consists of an intact endogenous *galK* gene of *E. coli* that harbors a random promoter sequence (P0) that replaces its endogenous promoter. By growing *E. coli* in the presence of the sugar galactose, we are selecting for increased *galK* expression. Adaptation to selection for increased expression can happen by two different, non-mutually exclusives ways: through increased copy-number (duplication or amplification) or through point mutations in the P0 promoter region of *galK* (divergence) (Tomanek *et al*., 2020).

Importantly, our genetic reporter system allows us to distinguish between the two mutation types. *GalK is* part of a chromosomal reporter gene cassette and is transcriptionally fused to a *yfp* gene (Fig. 1a). Hence, any increases in *galK* expression – be it by copy-number or point mutations – can be detected as increases in YFP expression. However, only mutations to the copy-number of the entire *galK* locus lead to an additional increase in the expression of an independently transcribed *cfp* gene downstream of *galK-yfp* (Steinrueck and Guet, 2017; Tomanek *et al*., 2020) (Fig. 1a). Hence, increases in *yfp* alone indicate the divergence of the *galK* promoter sequence P0 by point mutations, while increases of both fluorophores indicate copy-number mutations of the whole locus. Finally, clones with increased *yfp* but without point mutations in P0 would indicate the presence of a trans-acting mutation at a different locus on the chromosome or a rare amplification event occurring independent of the repeated IS elements and excluding CFP (Steinrueck and Guet, 2017; Tomanek *et al*., 2020). Moreover, while in principle possible, an adaptive mutation in the coding sequence of *galK* itself is extremely unlikely to be selected under our experimental conditions given that growth is limited only by expression of the endogenous and fully functional galactokinase enzyme.

**Figure 1.**
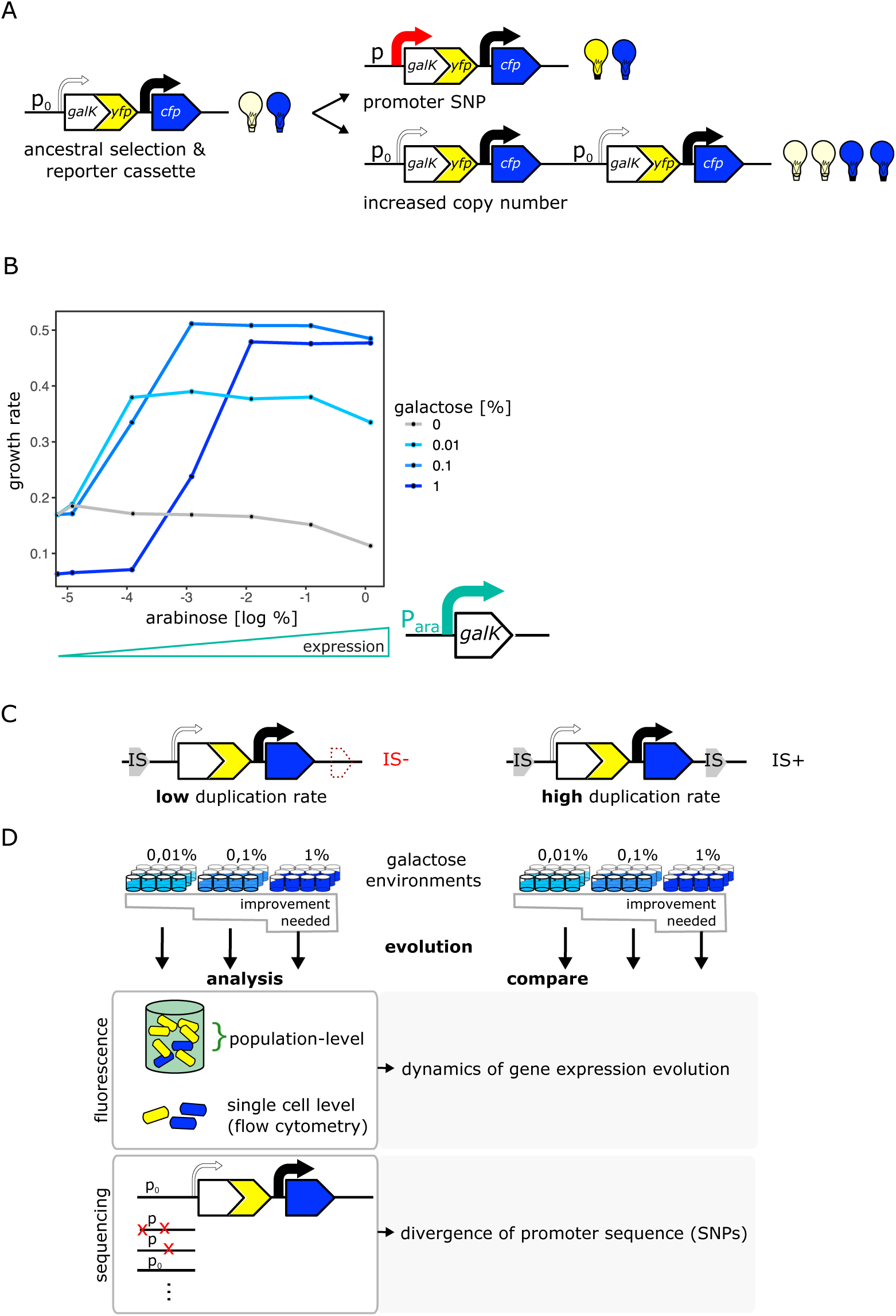
An experimental system to study gene duplication and divergence in strains with different duplication rates. **A**. Cartoon of chromosomal selection and reporter cassette. The *galK-yfp* gene fusion does not have a functional promoter, but instead a random sequence, P0 (thin arrow), drives very low levels of baseline gene expression. *Cfp* expression is driven by a constitutive promoter (black arrow). Light bulbs symbolize fluorescence. Two fundamentally different kinds of adaptive mutations are shown on the right: (i) point mutations in P0 lead to increases in GalK-YFP while CFP remains at ancestral single-copy levels (top), (ii) mutations to the copy number of the whole reporter cassette will increase both YFP and CFP expression (bottom). **B**. Growth rate (as a proxy for fitness) as a function of different induction levels of *galK* expression in four different concentrations of galactose. Expression of a synthetic p_ara_-*galK* cassette (schematic below the figure) is induced by the addition of arabinose. Growth rate increases along with increasing *galK* expression, but it plateaus at different values for different gene expression levels depending on galactose concentration (low, intermediate and high gene expression demand). **C-D**. Experimental layout. The adaptive dynamics and sequence divergence in P0 is compared between two otherwise isogenic strains (IS- and IS+) that differ in their rate of forming duplications. For IS-the second endogenous copy of *IS*1C located 12kb downstream of the selection and reporter cassette has been deleted (**C**). 96 replicate populations of each strain are evolved in three different levels of galactose, which select for increasing levels of gene expression improvement for twelve days, respectively. Throughout, fluorescence is analysed in bulk and on a single cell level to analyse evolutionary dynamics, and relevant clones are sequenced (**D**).

### Different substrate levels result in different enzyme expression demands

Our experimental environment consists of liquid minimal medium containing amino acids as a basic carbon and energy source, such that cells can grow even in the absence of *galK* expression (Fig. 1b – grey line). Adding galactose to this basic medium renders *galK* expression highly beneficial. To characterize the relation between fitness and *galK* expression, we engineered a construct where the expression of *galK* is induced by the addition of arabinose. Growth rate increased along with *galK* expression and saturated at a certain expression level, which depended on the galactose medium used (Fig. 1b). Thus, our system allows studying adaptation in environments with different gene expression demands: low concentrations of galactose demand a low level of *galK* expression (and increasing expression above this level does not add any extra benefit), while high concentrations of galactose demand a higher level of *galK* expression to obtain maximum growth rate. In other words, our experimental system allows selecting for different levels of improvement of a biological function (in our case increased *galK* expression) by growing cells in different galactose concentrations.

### Evolution of galK expression in IS+ and IS-strains

Given the vast range of duplication rates observed at different chromosomal loci in bacteria (Roth *et al*., 1988; Andersson and Hughes, 2009; Elez *et al*., 2010; Reams and Roth, 2015), our objective was to experimentally manipulate the ability of *galK* to form duplications and study its effect on evolutionary dynamics. A common way to manipulate the duplication rate is by deleting the *recA* gene involved in homologous recombination (Goldberg and Mekalanos, 1986; Reams *et al*., 2010; Dhar, Bergmiller and Wagner, 2014). However, given its role in DNA repair, comparing *recA* and *Λ 1recA* strains will be strongly influenced by the growth defects that such a mutation entails. In order to not have to consider pleiotropic effects caused by a difference in the genome-wide duplication rate, we instead compare two identical strains whose difference in duplication rate is restricted to a single genomic locus. To this end, we take advantage of a chromosomal location that is characterized by high rates of duplication and amplification due to homologous recombination occurring between two endogenous identical insertion sequences (IS) elements that flank this specific locus (Steinrueck and Guet, 2017; Tomanek *et al*., 2020). By deleting one copy of IS*1*, we generated two otherwise isogenic strains of *E. coli* that differ solely by the presence of one IS*1* element approximately 10 kb downstream of *galK* (Fig. 1c), and are thus predicted to show strong differences in their rates of duplication formation at this locus. In the following, we will refer to these strains as IS+ and IS-.

To understand how the duplication rate affects adaptive dynamics we conducted an evolution experiment with 96 replicate populations of the IS+ and IS-strains (Fig. 1 d). Growing these populations in minimal medium containing only amino acids (control) or supplemented with three different galactose concentrations enabled us to follow adaptation to different gene expression demands (levels of selective pressure) (Fig. 2a). Daily measurements of population fluorescence prior to dilution (1:820) allowed us to monitor population phenotypes roughly every ten generations over twelve days.

**Figure 2.**
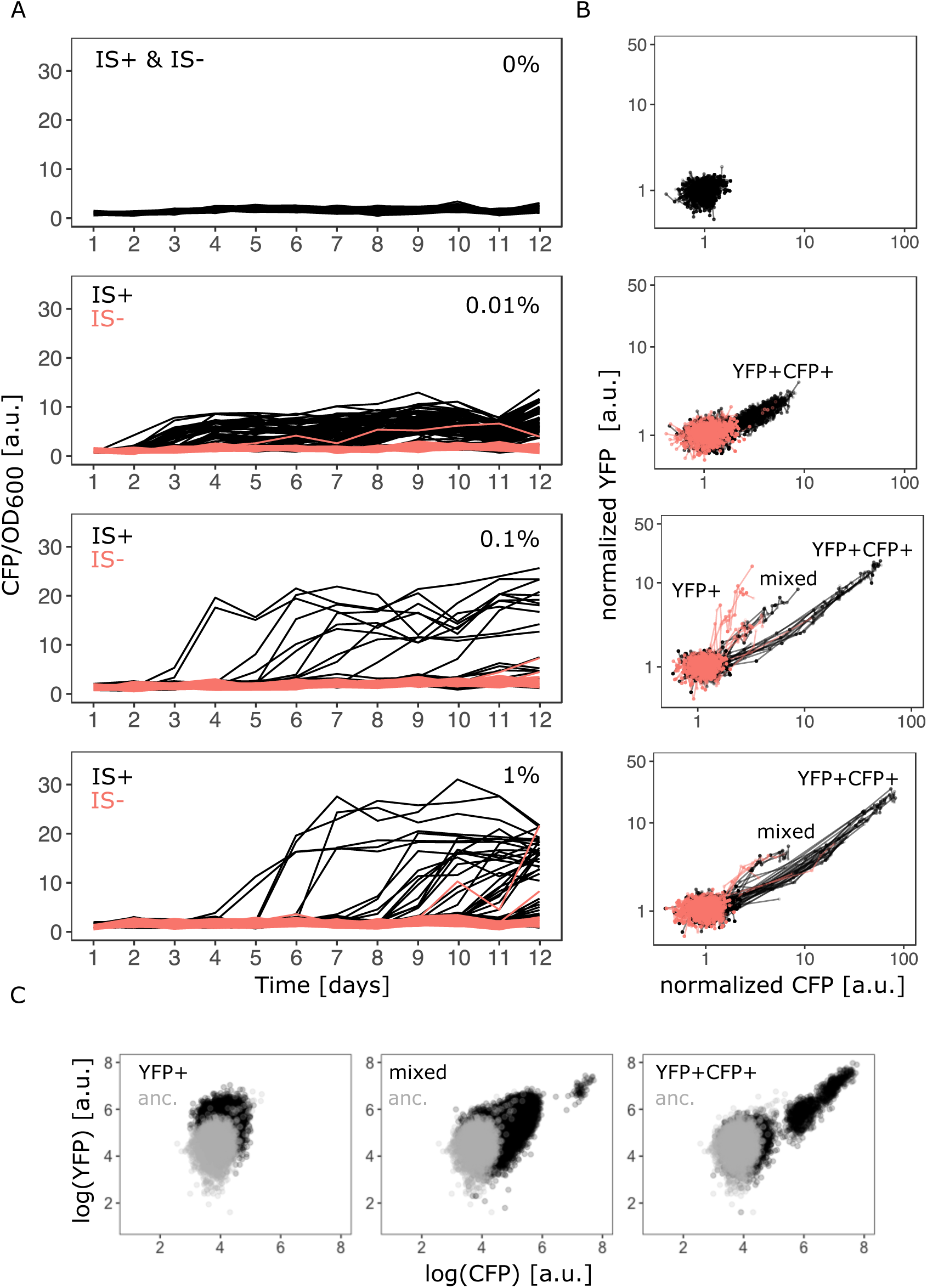
Evolutionary dynamics depend on galactose concentration and duplication rate. **A**. Daily measurements of normalized CFP fluorescence as a proxy for gene copy-number of 96 populations of IS+ (black) and IS- (red) strains growing in three different galactose concentrations (% indicated in the plot), respectively, as well as 33 replicates of IS+ and IS-strain, respectively, growing in the absence of galactose (control, black). **B**. Logarithmic plots for an overview of fold changes in YFP and CFP fluorescence of populations from (**A**) (YFP and CFP were normalized to the mean fluorescence of ancestral populations (Anc) evolved in 0% galactose (top panel)). Lines connect measurements of each population. Populations’ fluorescence phenotypes occupy three different areas: increased YFP only (YFP+), increased CFP and YFP (YFP+CFP+, i.e. amplified) and increased CFP with an additional elevation in YFP above the YFP+CFP+ fraction (mixed). **C**. Representative flow cytometry plots showing single-cell YFP and CFP fluorescence for populations from the YFP+ (left), mixed (middle) and YFP+CFP+ (right) fraction (indicated in panel **B**), respectively.

The evolution experiment confirmed that the two strains differ strongly in their rate of copy-number mutations of the *galK* locus. The strain lacking one of the flanking IS*1* elements (IS-) showed a drastic reduction in the ability to undergo *galK* amplification. In contrast to the IS+ strain, very few IS-populations evolved increased CFP expression (Fig. 2a – red traces). Interestingly, in the IS+ strain, the number of populations amplified by the end of the experiment depended on the environment. At least twice as many populations were amplified in the low (0.01%) galactose environment compared to the other two environments (68, 19 and 34 populations for low, intermediate and high galactose, respectively) (Supplementary Fig. 1a). Not only the number of amplified populations, but also the maximum CFP fluorescence attained by IS+ populations differed significantly between the low (0.01%) and higher (0.1% and 1%) galactose environments (Supplementary Fig. 1b). Populations, which evolved increases in CFP fluorescence did so within two days and maintained this level relatively stably for the duration of the experiment. (See Supplementary Fig. 2a for an independent evolution experiment confirming the environment-dependent patterns of amplification.) The observed difference in the number of *galK* copies is consistent with the observation that the three environments select for different levels of increasing gene expression (‘levels of improvement’) (Fig. 1b) and confirms that amplifications are an efficient way of tuning gene expression (Tomanek *et al*., 2020).

We then asked whether other differences in the nature of adaptive mutations exist between the three different environments. To get a coarse-grained overview, we plotted the YFP fluorescence of evolving populations as a proxy for *galK* expression against their CFP fluorescence as a proxy for *galK* copy-number for all time points (Fig. 2b). The YFP-CFP plot shows that evolving populations exhibit qualitatively different distributions of fluorescence levels in the three different environments, indicating that adaptation has followed different trajectories.

In the absence of galactose, populations retain their ancestral fluorescence phenotype. In the lowest galactose concentration (0.01%), data points show a correlated increase between YFP and CFP fluorescence indicative of gene-copy number mutations (“YFP+CFP+” in Fig. 2b). In the intermediate galactose concentration (0.1%) the IS-populations exhibit increased YFP fluorescence with ancestral (single-copy) CFP fluorescence indicative of promoter mutants, (“YFP+” fraction in Fig. 2b; Supplementary Fig. 3a). However, sequencing the P0 region upstream of *galK* of these evolved clones from populations with strongly increased YFP fluorescence (“YFP+” fraction in Fig. 2b) showed that they harbored an ancestral P0 sequence (Supplementary Fig. 3a). We hypothesized that the YFP+ populations carried an amplification extending into *galK*-*yfp*, yet excluding *cfp*. Quantitative real-time PCR confirmed our suspicion (Supplementary Fig. 3b). As the IS-strain cannot undergo the frequent duplication via the two flanking IS elements, it cannot access a major adaptive route available to the IS+ strain. Thus, its adaptation follows an alternative trajectory, which occurs through a repeat-independent lower-frequency duplication with junctions between *yfp* and *cfp* (Supplementary Fig. 3c). Despite the occurrence of *yfp*-only mutations in the IS-strain, increased CFP still reliably reports on increased copy-number. However, the *yfp*-only amplification hijacks our ability to unambiguously infer ancestral copy-number from ancestral CFP fluorescence alone. Instead, ancestral copy-number can only be confirmed by qPCR. However, we were ultimately interested in the divergence of promoter sequences, and going forward relied on sequencing to unambiguously determine the presence of adaptive promoter mutations.

In the high (1%) and intermediate (0.1%) galactose environment, data points occupy an additional space (“mixed fraction” in Fig. 2b) between the other two fractions, where both YFP and CFP are increased, but the YFP increase is larger than in the YFP+CFP+ fraction. Based on these population-level data, we hypothesized that this phenotypic space is occupied either by a population of mixed mutants carrying a combination of point and copy-number mutations, or by populations consisting of cells with only promoter mutations and cells with only copy-number mutations, (i.e. the two mutations being mutually exclusive). Knowing the single cell phenotype is therefore crucial for distinguishing between the two cases. Importantly, single cell fluorescence (using FACS) recapitulated the population measurements with the YFP-CFP phenotype falling into three distinct fractions (Fig. 2c).

### Copy-number and point mutations occur as a combination in the intermediate and high demand environment

To understand whether copy-number and point mutations are mutually exclusive or if they occur as a combination in the IS+ strain after evolution in intermediate (0.1%) and high (1%) galactose, we determined the single-cell fluorescence of all mixed fraction populations using flow cytometry (Figure 3a-b). It is worth noting that after twelve days of evolution, cells with ancestral YFP and CFP fluorescence were still present in every single amplified population. While some populations consisted of a high fraction of cells with elevated CFP fluorescence, mutants did not yet spread to complete fixation in any of them, highlighting the fact that our experiments are capturing the transient adaptive dynamics.

**Figure 3.**
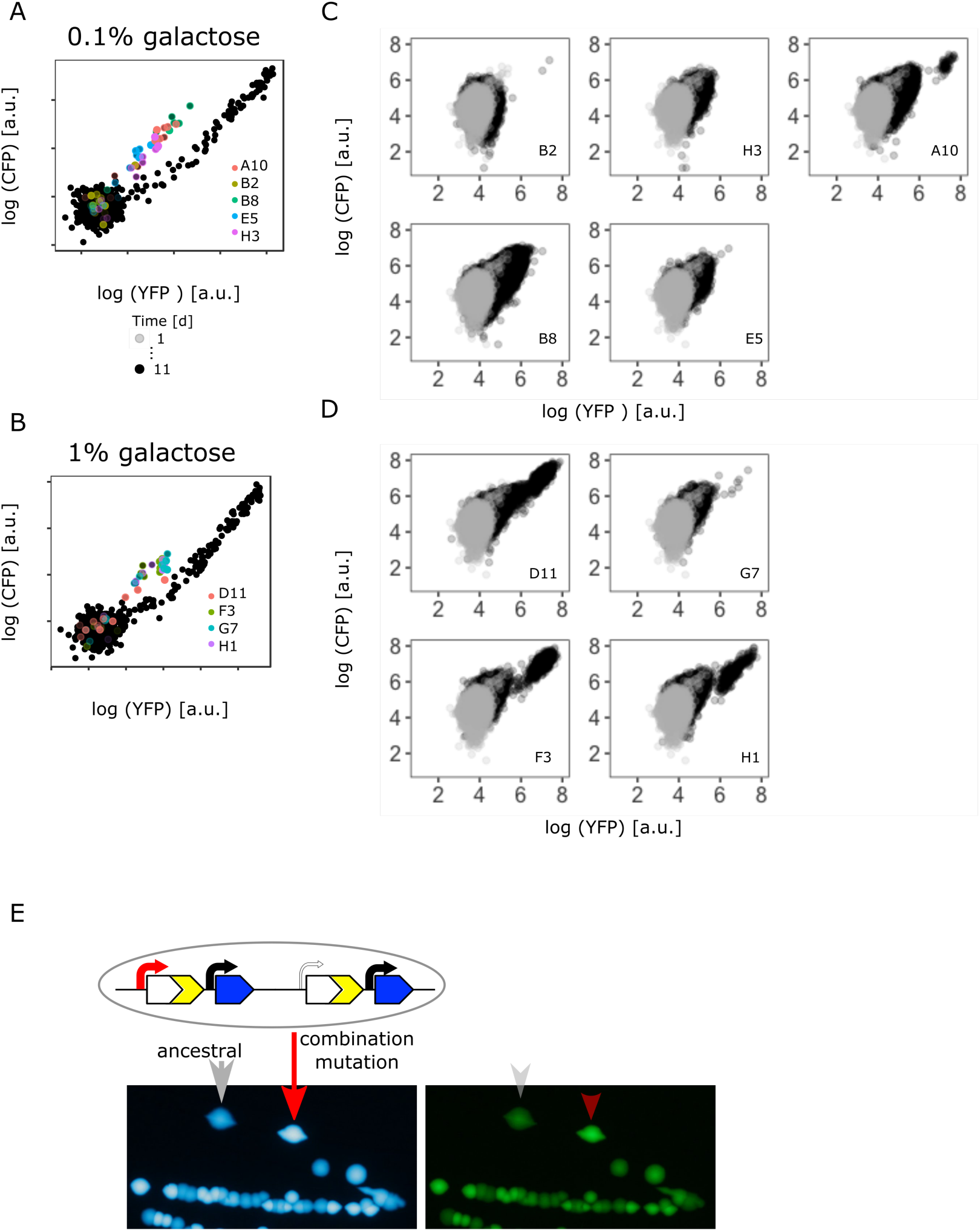
Confirming the presence of a combination of copy-number and point mutations in intermediate and high galactose. **A-B**. Log plot of YFP and CFP fluorescence of all 96 IS+ populations during evolution in 0.1% (A) and 1% (B) galactose (black points), respectively. Data replotted from Fig. 2B for an overview of population fluorescence of all mixed fraction populations (colored points). Time points of measurements are indicated by the degree of shading. **C-D**. Single cell fluorescence phenotypes as measured by flow cytometry of all mixed fraction populations identified in (**A-B**) after twelve days of evolution, respectively, indicate the presence of combination mutations (an increase of both YFP and CFP within a single cell as opposed to a mixed population of cells with either an increase in YFP or an increase in CFP, compare to Fig. 2C). **E**. Sanger sequencing of individual colonies allows to determine the genotype of an evolved clone of any fluorescence phenotype. Images of CFP (left) and YFP (right) fluorescence of individual colonies from a representative IS+ population (A10) streaked onto LB agar after having evolved in 0.1% galactose for twelve days. Sanger sequencing of the P0 sequence revealed a T>A point mutation in an amplified (red arrow) but not an ancestral colony (grey arrow).

Flow cytometry results showed that IS+ populations of the mixed fraction from intermediate (0.1%) galactose (Fig. 3a) consisted of a single type of mutant with increased YFP/CFP fluorescence relative to the ancestral values (Fig. 3c). If instead a population consisted of two mutually exclusive mutants, we would expect cells to fall into two distinct phenotypic clusters, one with only increased YFP (corresponding to the “YFP+” fraction) and one with only amplifications (corresponding to the “YFP+CFP+” fraction). Moreover, YFP fluorescence of the mixed fraction cells was greater than YFP for pure amplification mutants, which falls along the diagonal axis (Fig. 2c - right panel), again indicating a combination of copy-number and promoter mutations. To confirm the presence of combination mutants, we randomly picked three populations of the mixed fraction. Sequencing revealed that within these populations, only amplified clones, but not clones with single-copy *cfp* harbored a SNP (-30T>A) in P0 (Fig. 3e).

Similar to intermediate galactose, IS+ populations from the high (1%) galactose mixed fraction (3b) harbored cells with the combination mutation phenotype and, in addition, cells with pure amplifications (Fig. 3d). Taken together, these data indicate that copy-number and point mutations can occur as a combination in environments with sufficiently high gene expression demand.

### Copy-number and point mutations are mutually exclusive in the low demand environment

After finding combined mutants in the high galactose environments, we analyzed the single cell fluorescence of all IS+ populations from the low (0.01%) galactose environment. Suprisingly, and in contrast to the intermediate and high galactose environments, in low galactose adaptive amplification of IS+ populations happened rapidly with the majority of populations showing increases in CFP fluorescence during the course of the experiment (Fig. 4a – left top and bottom panel). Notably, cells of those few populations that did not follow this general trend (Fig. 4a – middle top and bottom panel) showed an increase in YFP without a concomitant increase in CFP. As this small increase in YFP was not visible in the initial population measurements of liquid cultures (Fig. 2b), we turned to patching populations onto LB agar, a potentially more sensitive method which alleviates changes in fluorescence related to growth-rate. Imaging populations confirmed the increase in YFP for all populations with elevated YFP in single cell measurements (Fig. 4b & d). We examined individual populations with clearly increased YFP levels more carefully by re-streaking them on LB agar (Fig. 4c). Consistent with flow cytometry results (Fig. 4a - right panel), we found colonies with three different fluorescence phenotypes: ancestral, increased YFP (“YFP+”), and a small subpopulation with both, increased YFP and CFP (amplified). Sequencing of the amplified colony type confirmed it to be a *bona fide* amplification without additional promoter SNPs. Sequencing of the YFP+ colony uncovered two adaptive SNPs in P0 (-30T>A and -37C>T), which were identical to a previously identified promoter mutation “H5” (Supplementary Fig. 2b) (Steinrueck and Guet, 2017; Tomanek *et al*., 2020).

**Figure 4.**
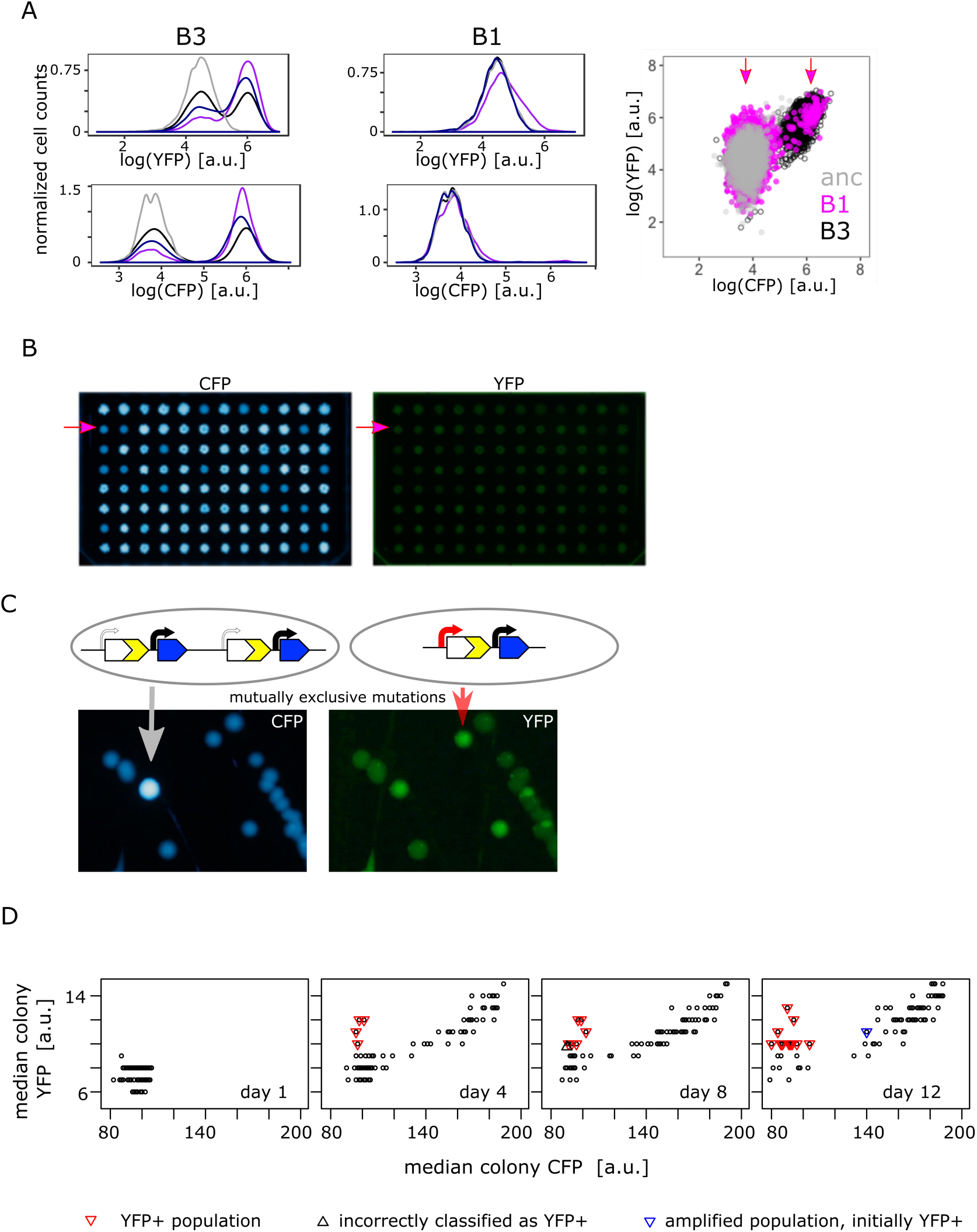
Confirming the presence of mutually exclusive mutations in low galactose. **A**. Representative flow cytometry histogram showing YFP fluorescence (upper left and middle panel) and CFP fluorescence (lower left and middle panel) of IS+ populations B3 (left panels) and B1 (middle panels) over time (grey - ancestral, black - day 4, dark blue - day 8, purple – day 12). Right panel shows the same data for populations B3 and B1 as a YFP versus CFP plot in order to better visualize the two distinct sub-populations in B1 (magenta). **B**. Representative images of CFP (left panel) and YFP (middle panel) fluorescence of populations patched onto LB agar, which allows comparing population fluorescence in the absence of galactose-dependent growth effects. Magenta arrows indicate population B1, which exhibits increased YFP but ancestral CFP fluorescence (quantification of patch fluorescence intensity in **D**). **C**. Images of CFP (left) and YFP (right) fluorescence of individual colonies from IS+ population B1 (shown in **B**) streaked onto LB agar after twelve days of evolution in 0.01% galactose. The population consists of amplified colonies with increased CFP and YFP fluorescence (grey arrows) and single-copy colonies with a promoter mutation (red arrows). **D**. Quantitative analysis of patched populations indicates that promoter mutants (YFP+) evolve only in single-copy backgrounds. YFP-CFP plot of median colony fluorescence intensity of populations patched onto agar (as shown in (**B**)) on day 1, 4, 8 and 12 of evolution in 0.01% galactose. Populations were classified as YFP+ if their YFP but not CFP fluorescence intensity values exceeded ancestral fluorescence (red triangles, confirmed by flow cytometry). In all these populations, the YFP+ phenotype evolved from an ancestral phenotype. Blue triangle represents an amplified population, which was classified as YFP+ in the previous time point (flow cytometry showed that this population became dominated by copy-number mutations later). Grey triangle marks population incorrectly classified as YFP+ (ancestral fluorescence according to flow cytometry). See also Supplementary Table 1.

As we failed to find combination mutants (i.e. a mixed fraction) in population measurements from the low galactose environment (Fig. 2b), we used agar patches from four different time points of the evolution experiment to screen IS+ populations more comprehensively (Fig. 4d). Re-streaking, sequencing and flow cytometry analysis revealed that all populations with elevated YFP and ancestral CFP harbored either only promoter mutants or a mixed population of a few amplified cells and a majority of promoter mutants (Supplementary Table 1). As opposed to high and intermediate galactose, we did not find a single population with combined mutants in low galactose. Moreover, the fact that mutations were mutually exclusive within populations, was also reflected when we analyzed their fate over time. Quantitative analysis of the fluorescence intensity of patched populations (Fig. 4d), confirmed that populations with a significant fraction of promoter mutants (i.e. visibly YFP+ on the agar patch) did not become amplified later in the experiment. As a single exception, population F6 gained the YFP+ phenotype early, but became dominated by gene amplifications by the end of the experiment (Fig. 4d –right panel, blue triangle). Nevertheless, also in this case, copy-number and point mutations did not occur in the same genetic background. Conversely, all YFP+ populations evolved exclusively from those with ancestral phenotype; no single amplified population gained a functional promoter within the time frame of the experiment (Fig. 4d).

The complete absence of combined mutants in the low demand environment is consistent with the fact that only a modest increase in *galK* expression is necessary to reach maximal fitness (Fig.1b). Thus, while a combination of amplification and promoter point mutation evolves in response to selection for a strong increase in *galK* expression (intermediate and high demand environments), either mutation alone might provide a sufficient increase in gene expression to allow for maximal growth in the low demand environment. This means that the fitness benefit of either mutation does not add up when combined. In other words, negative epistasis precludes the evolution of combination mutants in the low demand environment.

### An increased fraction of adaptive promoter mutations is found in IS-populations evolved in the low demand environment

If point mutations are more frequent than copy-number mutations and do not occur as a combination in the low demand environment, we would expect divergence to proceed more slowly as compared to an intermediate or high demand environment.

To directly test this hypothesis, we estimated the level of divergence between all of the IS+ and IS-populations evolved in the low demand (0.01% galactose) environment. We pooled all 96 populations into pools of 32 and quantified the fraction of SNPs in P0 previously known to be adaptive (Tomanek *et al*., 2020). To do so, we subjected PCR amplicons of the pooled populations to next generation sequencing (Fig. 5a, Supplementary Fig. 4a). We designed our sequencing experiment such that we were able to analyze 39bp upstream and downstream of the *galK* start codon. We counted the number of sequence reads carrying either one or both most frequently observed adaptive SNPs at position -30 and -37 upstream of the *galK* start codon (Supplementary Table 1). As a control, we also compared the number of SNPs within the *galK* gene of the IS+ and IS-evolved under different galactose conditions. In our experimental system, galactose-selection is not expected to lead to adaptive mutations anywhere in the coding region of *galK*, as the enzyme itself is fully functional despite lacking a functional promoter sequence. As the absolute number of sequencing reads differs for each sample (Supplementary Fig. 4a), a meaningful comparison of the number of SNPs between different environments can only be achieved by normalizing to the respective number of ancestral reads of each sample. We therefore counted the number of sequencing reads with either zero mismatches (ancestral sequence) or one single mismatch (single SNP). Consistent with our expectation, the mean number of sequencing reads with a single SNP at any position in *galK* was similar in populations evolved in different galactose concentrations and in the control populations evolved in the absence of galactose (Supplementary Fig. 4b). We then compared the fraction of reads with the two adaptive SNPs in P0 previously known to confer increased *galK* expression (Fig. 5a). While the fraction of reads carrying SNPs in *galK* is similar in all media, SNPs in P0 were more frequent in media containing galactose than in the control (Fig. 5a left and right panel) in agreement with strains adapting to galactose selection. Intriguingly, in low galactose, we found a higher fraction of reads carrying both adaptive single SNPs (-30T>A and - 37C>T) in IS-populations than in the IS+ populations. This is consistent with our hypothesis that the more frequent amplification mutants effectively out-compete point mutations in the low demand environment.

**Figure 5.**
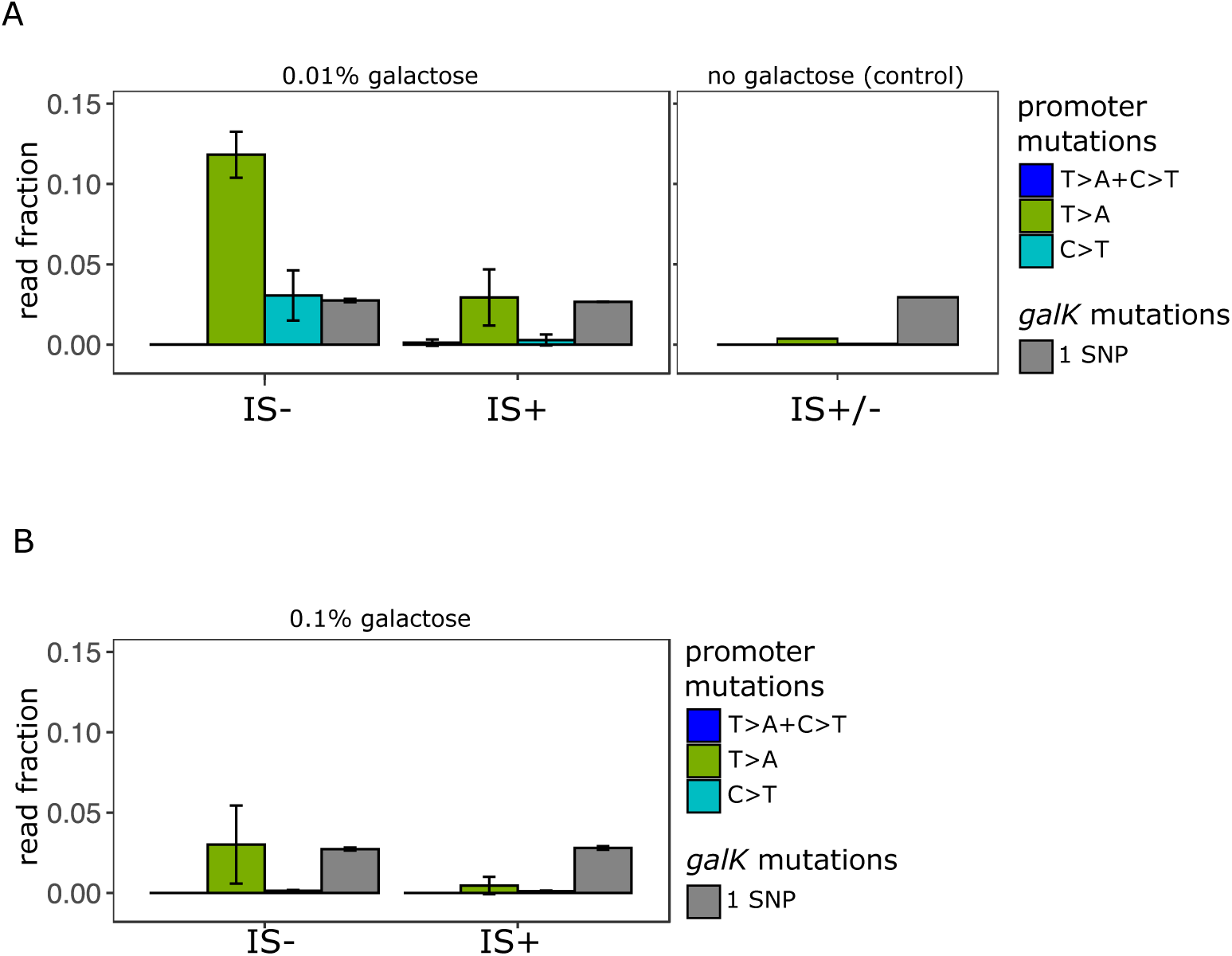
Amplicon deep sequencing of P0 in pooled evolved populations. **A**. (Left panel) Number of reads carrying a P0 sequence with two adaptive SNPs 30 bp and 37 bp upstream of *galK*, respectively, (“T>A+C>T” in blue) or its respective single SNPs (“T>A” in green, “C>T” in cyan). Values are normalized to the number of reads with ancestral P0 for IS- and IS+ populations evolved in 0.01% galactose. The mean fraction of reads with any single SNP in *galK* is shown as a control (grey). Error bars represent the standard deviation of three replicates, consisting each of 32 pooled evolved populations. (Right panel) Read fractions of the same respective SNPs shown for a pool of all 96 IS+ and IS-populations evolved in the absence of galactose. **B**. Mean read fractions as in (**A**) shown for three replicates of each 32 pooled populations evolved in intermediate (0.1%) galactose.

We are here using the fraction of sequencing reads (“alleles”) with adaptive SNPs divided by the total number of reads as a simple metric of divergence. However, this normalization, leads to an “underestimation” of SNPs if they occur in an amplified background. For instance, a SNP within a cell with four P0-*galK* copies, where one carries a SNP, counts less than a cell with one copy of P0-*galK* carrying one SNP. The rationale for using the fraction of adaptive alleles as our metric of divergence as opposed to the alternative, which is the number of SNPs per cell, is twofold: First, the methodology used here does not allow comparing absolute read counts between samples. Second, and more importantly, due to the random nature of deletion mutations, a single SNP in an amplified array of four copies has a 1 in 4 chance of being retained as a lasting divergent copy in the process of amplification and divergence. Hence, the “dilution” of SNPs by additional amplified copies is not simply a counting artifact, but reflects a biological reality relevant to the very process that we are studying. Therefore, we conclude that in the low demand environment a strain which cannot adapt by gene amplification exhibits a higher level of divergence than a strain which frequently adapts by gene amplification.

### Evolutionary dynamics between mutation types differ for different initial random promoter sequences

Given the paucity of point mutations that we observed for the evolution of the random P0 sequence (either a combination of -30T>A and -37C>T or each SNP alone), we wondered whether a greater variety of mutations could be obtained when using a different random promoter sequence as a starting point for evolution. Therefore, we repeated our evolution experiment in the intermediate (0.1%) galactose environment with three additional random promoter sequences (P0-1, P0-2, P0-3).

After ten days of evolution, only two out of the four random P0 sequences evolved increased *galK-yfp* expression (Fig. 6a). This is roughly consistent with the fact that approximately 60%-80% of random sequences are one point mutation away from a functional constitutive promoter (Yona, Alm and Gore, 2018; Lagator *et al*., 2020). Interestingly, P0-1 and P0-3 did not gain any gene duplications or amplifications. At first glance, this drastic difference in gene amplification was unexpected, since the IS+ strains only differ in their P0 sequence, and not in their gene duplication rate. However, random sequences have different abilities to recruit RNA-polymerase, and as a result, different baseline expression levels (Yona, Alm and Gore, 2018; Lagator *et al*., 2020). Given that a plateau exists in the expression-growth relation for low levels of expression (Fig. 1c), the initial expression level conferred by P0-1 and P0-3 might be too low to yield a selective benefit upon gene duplication alone. According to this hypothesis, these random (non-)promoters are not only two (or more) point mutations away from a beneficial sequence, but also two (or more) copy-number mutations.

**Figure 6.**
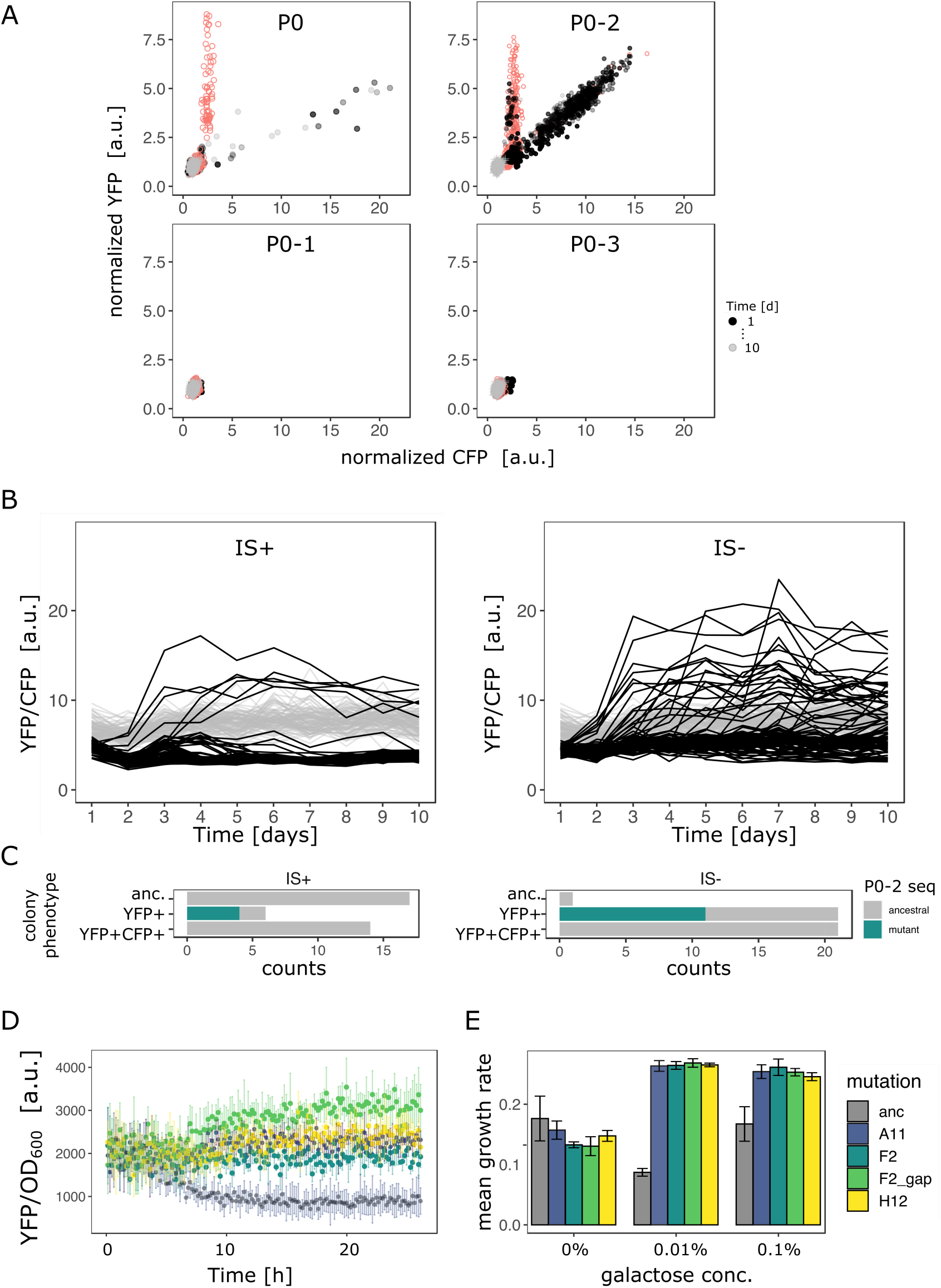
Evolutionary dynamics for different random P0 sequences in 0.1% galactose. **A**. YFP versus CFP fluorescence normalized to the ancestral value of 96 populations of IS+ (black) and IS- (red) strain each harboring a different random sequence upstream of *galK* (“P0”, “P0-1”, “P0-2”, “P0-3”) grown in 0.1% galactose and without galactose (grey lines, control), respectively. Time points are indicated by the degree of shading. **B**. YFP/CFP fluorescence to visualize increases in *galK-YFP* expression not caused by copy-number increases plotted for the duration of the evolution experiment for P0-2 populations of IS+ (left panel) and IS- (right panel). Here, gene amplifications are visible as slight decrease in YFP/CFP (see also Supplementary Fig. 5A) relative to the 0% galactose control (grey), putative promoter mutations are visible as an increase in YFP/CFP. **C**. Distribution of P0-2 mutants in IS+ and IS-populations after twelve days of evolution in 0.1% galactose. Mutations in P0-2 are exclusively found in populations with increased YFP and ancestral CFP fluorescence (YFP+). IS+ clones from all six YFP+ populations were sequenced, while IS-clones from a random subsample of 21 YFP+ populations were sequenced. **D**. Mean normalized YFP fluorescence of reconstituted P0-2 mutants and the P0-2 ancestor strain (grey) grown in control medium (0% galactose). **E**. Mean growth rate of reconstituted P0-2 mutants and the ancestor strain (grey) in 0.01% galactose, 0.1% galactose and control medium (0% galactose). Error bars represent the standard deviation of four replicates.

### Copy-number and point mutations are mutually exclusive in the intermediate demand environment for P0-2

For P0, the evolution experiment in intermediate galactose reproduced our previous findings, namely a YFP+CFP+ (amplified) and a mixed (amplified with increased YFP) fraction for IS+ populations and a YFP+ fraction for IS-populations (compare Fig. 6a with Fig. 2b), which corresponds to an amplification of YFP, but not CFP (Supplementary Table 2).

For P0-2, the evolutionary dynamics differed from P0. In the IS+ strain, almost every single population evolved amplifications within the first two days of the evolution experiment (Fig. 6b, Supplementary Fig. 5). Moreover, only two fractions are visible in the YFP-CFP plots of P0-2. The first fraction is occupied by YFP+ populations carrying a single copy of *cfp*. The second fraction along the diagonal between YFP and CFP, is occupied by amplified populations (YFP+CFP+). Moreover, it is shifted towards higher values of YFP/CFP relative to values found for P0, suggesting that P0-2 exhibits a higher baseline expression level than all the other three random promoter sequences. In contrast to the population-level measurements, single cell measurements were not sufficiently sensitive to corroborate any difference in leaky expression amongst the four random promoter sequences (Supplementary Fig. 5b). However, in line with the observed evolutionary dynamics, P0 and even more so P0-2 confers a significant growth advantage over the other two promoters (Supplementary Fig. 5c). As mentioned above, this suggests that the observed growth advantage of P0-2 populations can explain their rapid amplification dynamics. In agreement with the evolution experiments with P0, the YFP+CFP+ (amplification) fraction is also strongly reduced in the IS-strain for P0-2.

Intriguingly, with the majority of P0-2 IS+ population amplified, those few P0-2 IS+ populations that failed to evolve amplifications show an increase in YFP/CFP early in the evolution experiment (Fig. 6b – left panel). This result combined with the idea that P0-2 exhibits a relatively high baseline expression level and the absence of a mixed fraction for P0-2 (Fig. 6a), suggests that increases in gene expression evolve *either* via gene amplification *or* via point mutation. In other words, because initial *galK* expression is high in P0-2, a small improvement (either amplification or a promoter mutation) is sufficient to reach the required gene expression demand. Thus, the adaptive trajectory of P0-2 in intermediate galactose resembles that of P0 in low galactose as both environments select only for a modest improvement in *galK* expression.

In contrast to the IS+ strain, where only six populations showed increased YFP/CFP fluorescence that emerged only within the first three days of evolution, populations of the IS-strain were evolving increased YFP/CFP fluorescence throughout the experiment (Fig. 6b – right panel). We were curious whether the increase in YFP/CFP in both, IS+ and IS-populations, was due to promoter mutations. Sequencing of randomly picked evolved clones revealed that in the majority (4/6 for IS+, 11/21 for IS-) of clones with increased YFP/CFP indeed harbored a mutation in P0-2, including a SNP, a 12bp and a 13bp deletion (Supplementary Table 2; Figure 6c).

Importantly, colonies of the same populations but with ancestral fluorescence harbored ancestral P0-2 sequences (Table 1), indicating that the observed mutations (Supplementary Table 2) are causal for the increased YFP expression. While finding the causal mutations for the remaining evolved clones with increased YFP but ancestral P0-2 (Fig. 6c) lies outside the scope of the current work, we speculate that they may occur further upstream of P0-2 or could be acting *in trans* such as loss of function mutants in the transcription factor *rho* (Steinrueck and Guet, 2017).

To confirm that the 12bp deletion mutation, the 13bp deletion mutation and the SNP were in fact adaptive, we reconstituted these mutations into the ancestral P0-2 strain, where they conferred increased YFP expression (Fig. 6d) resulting in increased growth in medium supplemented with galactose (Fig. 6e). The finding that the promoter mutations were responsible for increased *galK*-*yfp* expression was corroborated by the fact that these mutations occurred exclusively in populations with increased YFP but ancestral CFP, and were completely absent in amplified and ancestral colonies from a random set of 17 IS+ populations (Fig. 6c). It is worth noting that mutations observed in P0-2 were more diverse than those observed in P0 (seven different mutations including indels, an IS insertion and a SNP in P0-2 versus three different SNPs in P0 – compare Supplementary Tables 1 and 2). Thus, amplification can interfere with divergence not only by point mutations but also small insertions and deletions.

Taken together, the facts that i) the majority of IS+ populations become rapidly amplified, ii) with few promoter mutations arising exclusively in the first day in non-amplified populations (mutations are mutually exclusive) and iii) many more promoter mutations occur in IS-populations throughout the evolution experiment, strongly suggest that negative epistasis between frequent copy-number mutation and point mutations hinder fixation of the latter.

### Amplification hinders divergence by point mutations in the low demand environment

Taken together, our results suggest that the evolutionary dynamics of duplication/amplification and divergence depend on the level of gene expression increase selected for (Fig.7). In both environments, promoter point mutations evolve at a low rate directly in a single copy background. However, if rates of copy-number mutation are high, evolutionary dynamics are dominated by amplification. Irrespective of the environment, this amplification increases the mutational target size for rarer adaptive point mutations to occur. However, only if a strong increase in *galK* expression is selected for (high demand environment) the beneficial effects of both types of mutation add up, and we observe a combination of amplifications and point mutations to occur, in agreement with the IAD model (Bergthorsson, Andersson and Roth, 2007; Näsvall et al., 2012; Andersson et al., 2015) (Fig. 7a). In contrast, if only a modest level of gene expression increase is selected (low demand environment) (Fig. 1b), a single mutational event may be sufficient to provide it. Therefore, adaptation is dominated by the more frequent type of mutation, namely copy-number. In other words, amplifications effectively hinder divergence in the low demand environment due to their negative epistatic interaction with point mutations and we call this effect the Amplification Hindrance hypothesis.

**Figure 7.**
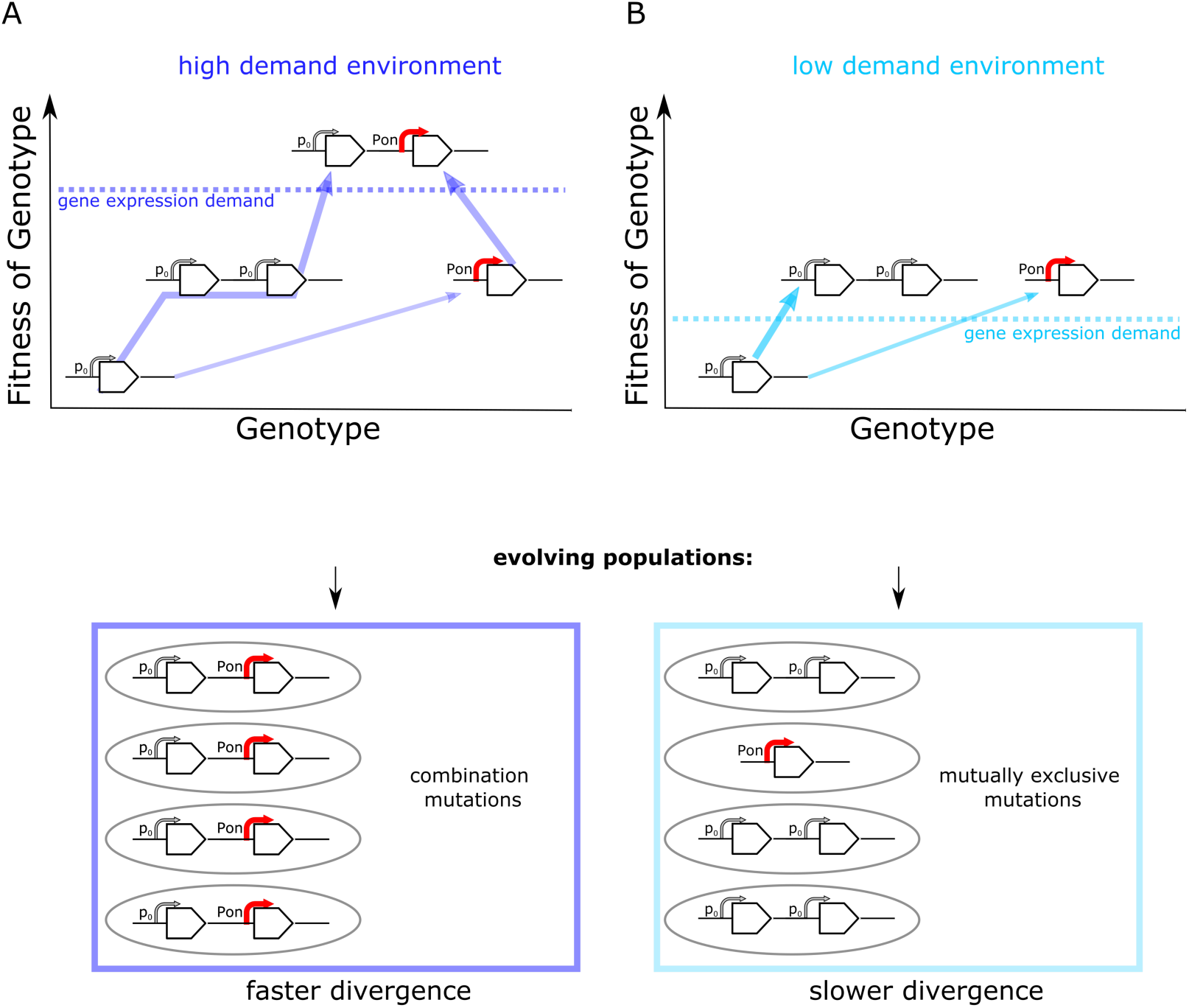
Frequent copy-number mutation can hinder adaptation by point mutations. Genotype-fitness map (‘fitness landscape’) illustrating the difference between adaptive trajectories of a high demand (**A**) and low demand (**B**) environment, which differ solely by the increase in gene expression they select for. The dashed line indicates the level of gene expression sufficient to reach maximal growth rate (‘fitness’) (see also Fig. 1B). Lower panels show the experimentally observed genotypes for each environment. **A**. For an environment selecting for a large increase in gene expression (high demand) more than one adaptive mutation is necessary to reach maximal fitness. If copy-number mutations are frequent (as in the IS+ strain), adaptation by amplification is most likely (bold arrow). Alternatively, at a lower frequency, adaptation occurs via a point mutation in the promoter sequence (thin arrow). Due to an increased mutational target size, cells with gene amplfications are more likely to gain a beneficial point mutation than cells with a single copy of *galK*. Alternatively, rare promoter mutants can become amplified, in either case leading to the combination mutant observed in experiments. **B**. For an environment selecting for only a modest increase in gene expression (low demand) maximal growth rate is attained either by gene amplification (more frequent, bold arrow) or point mutations (less frequent, thin arrow). Combination mutations are therefore not observed in the experiment.

## Discussion

In this study, we investigated the interaction dynamics between two different types of mutations, adaptive copy-number and point mutations. While the process of gene duplication and divergence *per se* has been intensely studied since the pioneering work of Ohno more than half a century ago, no experiments have scrutinized the early phase of this process, where transient evolutionary changes may prevail. So far, the few existing experimental studies simply introduced mutations *a priori* without studying their formation dynamics (Dhar, Bergmiller and Wagner, 2014), while *in silico* studies used genomics to query the ‘archeological’ results of millions of years of sequence evolution (Innan and Kondrashov, 2010).

Here we used experimental evolution to investigate how the early adaptive dynamics of diverging promoter sequences is influenced by the rate of copy-number mutations as well as the level of expression increase selected for. We found that the spectrum of adaptive mutations differed drastically between environments selecting for different levels of expression of the same gene (Fig.1b, 3a, 6a). Combined mutants carrying both, copy-number and promoter point mutations, only evolved under conditions selecting for big increases in the levels of *galK* expression. In contrast, selection for only a modest increase in *galK* expression lead to populations adapting by either gene amplifications or point mutations in their random promoter sequence, but not both simultaneously. Moreover, if amplification occurred early in the experiment, the random promoter sequence P0 did not diverge within the timespan of the experiment (Fig. 4d). This phenomenon was even more pronounced for a second random promoter sequence, P0-2 (Fig. 6b-c).

Moreover, comparing the number of point mutations between strains that differ solely in the rate of undergoing copy-number mutations in the *galK* locus, we found that under a low demand environment, a strain with a high duplication rate (IS+) diverged more slowly compared to a strain with low duplication rate (IS-).

Taken together, our results suggest that frequent gene amplification hinders the fixation of adaptive point mutations due to negative epistasis between these two different mutation types. While epistatic interactions can occur with any two adaptive mutations, copy-number mutations are unique, in that they are orders of magnitude more frequent than point mutations in bacteria (Roth *et al*., 1988; Drake *et al*., 1998; Andersson and Hughes, 2009; Elez *et al*., 2010; Reams and Roth, 2015) and in eukaryotes (Lynch *et al*., 2008; Lipinski *et al*., 2011; Schrider *et al*., 2013; Keith *et al*., 2016). This large difference in rates means that a competition between point and copy-number mutations is heavily skewed in favor of the latter (Figure 7b).

Unlike the phenomenon of clonal interference (which occurs between any two beneficial mutations even if their adaptive benefits are additive) (Gerrish and Lenski, 1998), negative epistasis does not slow down adaptation per se, as adaption is agnostic to whether point or copy-number mutations lead to an improved phenotype. However, negative epistasis slows down divergence as populations have reached the fitness peak with an alternative kind of adaptive mutation. Negative epistasis between point and copy-number mutations can be expected to occur in any selective condition, which requires only a relatively modest increase to a particular biological function, namely an increase in gene expression or enzyme activity by only a *few*-fold. Thus, Amplification Hindrance may not only be of general relevance for the evolution of gene expression in bacteria, but also for the evolution of promiscuous enzyme functions, which analogous to a barely expressed gene can be enhanced by either copy-number mutations or point mutations in the coding sequence.

While we found that amplification slows down divergence under conditions of negative epistasis, the consensus in the literature has been that copy-number mutations not only serve as a first step in the “relay race of adaptation” (Yona, Frumkin and Pilpel, 2015), but that they also facilitate divergence, either indirectly by providing a first “crude” adaptation to cope with a new environment until more refined adaptation occurs by point mutations, or directly by increasing the target size for point mutations (Andersson and Hughes, 2009; Elde *et al*., 2012; Yona, Frumkin and Pilpel, 2015; Cone *et al*., 2017; Bayer, Brennan and Geballe, 2018; Lauer *et al*., 2018; Todd and Selmecki, 2020). The intuitive idea that amplification speeds up divergence (Andersson, Slechta and Roth, 1998) was originally developed as strong evidence against the adaptive mutagenesis hypothesis proposed by Cairns and others (Cairns, Overbaugh and Miller, 1988; Cairns and Foster, 1991).

Based on it, various experimental studies interpreted observations of adaptation to dosage selection in the light of “amplification as a facilitator of divergence” (Song *et al*., 2009; Pränting and Andersson, 2011; Elde *et al*., 2012; Näsvall *et al*., 2012; Yona *et al*., 2012; Yona, Frumkin and Pilpel, 2015; Cone *et al*., 2017; Bayer, Brennan and Geballe, 2018; Lauer *et al*., 2018; Todd and Selmecki, 2020). However, despite showing that adaptive amplification *precedes* divergence by point mutations, none of the studies provided a direct experimental test of the hypothesis that amplification *causes* increased rates of divergence. Experiments controlling for the rate of amplification were needed in order to dissect the ensuing evolutionary dynamics and establish causality.

All else being equal, more copies indeed mean more DNA targets for point mutations to occur (San Millan *et al*., 2017). However, as our experiments show, all else is not necessarily equal, and the evolutionary dynamics may differ strongly between an organism that can increase copy-number as an adaptation and an organism that cannot. Intriguingly, indications for more complex dynamics can be found in the existing literature (Yona *et al*., 2012; Lauer *et al*., 2018; Richts *et al*., 2021). One study showed that rapid adaptive gene amplification in yeast results in strong clonal interference between lineages (Lauer *et al*., 2018). A second study in yeast found that adaptation to an abrupt increase in temperature was dominated by rapid copy-number mutation, with SNPs occurring only much later (Yona *et al*., 2012; Yona, Frumkin and Pilpel, 2015). In a third experimental evolution study adaptation was dominated by copy-number mutations and the authors noted the surprising lack of promoter mutations (Richts *et al*., 2021).

The transient dynamics of gene amplification allows tuning of gene expression on short evolutionary timescales in the absence of an evolved promoter (Tomanek *et al*., 2020). In principle, such transient evolutionary dynamics do not leave traces in the record of genomic sequence data on evolutionary time scales and as such, their detailed study may not seem warranted. This is especially true in the context of duplication and divergence of paralogs, which is studied because abundant genomic sequence data are available (Kondrashov, 2012). Our present study proved this intuition wrong, as we uncovered a potentially long-lasting effect resulting from the transient dynamics associated with copy-number mutations: if adaptation by amplification is the fastest and sufficient, other, less frequent, mutations may not have a chance to compete. However, adaptive amplification returns to the ancestral single copy state in the absence of selection. This means that once the selective benefit of the transient adaptation is over, no change at the level of genomic DNA remains (Roth *et al*., 1996). Therefore, the idea that gene amplifications act as a transient “regulatory state” rather than a mutation (Roth *et al*., 1996; Tomanek *et al*., 2020) can be extended by an implication found here, namely that amplifications could effectively act as buffer against long-lasting point mutations. Thus, on sufficiently long time-scales, the transient dynamics that play out before the fixation of mutations may ultimately shape entire genomes (Cvijović, Nguyen Ba and Desai, 2018). Finally, Amplification Hindrance is in agreement with the observation that duplication and divergence is not a dominant force in the expansion of protein families in bacteria (Treangen and Rocha, 2011; Tria and Martin, 2021). In some sense then, in all situations where rapid amplification provides sufficient adaptation, it could act as a passive mutational force that – in addition to purifying selection – acts to conserve existing genes and their expression level.

## Acknowledgements

We are grateful to N. Barton, F. Kondrashov, M. Lagator, M. Pleska, R. Roemhild and G. Tkacik for input on the manuscript and to K. Tomasek for help with flow cytometry. I.T. is recipient of an OMV PhD fellowship.

## Methods

### Bacterial strain construction

To construct the IS-strain, we replaced the second copy of IS*1* downstream of the selection and reporter cassette in IT030 (Tomanek *et al*., 2020) with a kanamycin cassette using pSIM6-mediated recombineering (Datta, Costantino and Court, 2006). Recombinants were selected on 25 µg/ml kanamycin to ensure single-copy integration.

To generate the additional random promoters sequences P0-1, P0-2 and P0-3, we generated 189 nucleotides using the “Random DNA sequence generator” (https://faculty.ucr.edu/~mmaduro/random.htm) with the same GC content as P0 (55%). We synthesized these three sequences as gBlocks (Integrated DNA Technology, BVBA, Leuven, Belgium) with attached XmaI and XhoI restriction sites, which we used to clone P0-1, P0-2 and P0-3 into plasmid pMS6* (Tomanek *et al*., 2020) by replacing P0. We used pMS6* with the respective P0 sequence as a template to amplify the selection and reporter cassette and integrate it into MS022 (IS+) and IT049 (IS-) as described previously (Tomanek *et al*., 2020).

>P0

ACCGGAAAGACGGGCTTCAAAGCAACCTGACCACGGTTGCGCGTCCGTATCAAGATCCTCTTAATAAGCCCCCGTCACTGTTGGTTGTAGAGCCCAGGACGGGTTGGCCAGATGTGCGACTATATCGCTTAGTGGCTCTTGGGCCGCGGTGCGTTACCTTGCAGGAATTGAGGCCGTCCGTTAATTTCC

>P0_1

GTAGGCCCGCACGCAAGACAAACTGCTGGGGAACCGCGTTTCCACGACCGGTGCACGATTT AACTTCGCCGACGTGACGACATTCCAGGCAGTGCCTCCGCCGCCGGACCCCCCTCGTGATC GGGTAGCTGGGCATGCCCTTGTGAGATATAACGAGAGCCTGCCTGTCTAATGATCTCACGG CGAAAG

>P0_2

TCGGGGGGACAGCAGCGGCTGCAGACATTATACCGCAACAACACCAAGGTGAGATAACTCCGTAGTTGACTACGCGTCCCTCTAGGCCTTACTTGACCGGATACAGTGTCTTTGACACGTTTGTGGGCTACAGCAATCACATCCAAGGCTGGCTATGCACGAAGCAACTCTTGGGTGTTAGAATGTTGA

>P0_3

CCCCTGTATTTGGGATGCGGGTAGTAGATGAGCGCAGGGACTCCGAGGTCAAGTACACCACCCTCTCGTAGGGGGCGTTCCAGATCACGTTACCACCATACCATTCGAGCATGGCACCATCTCCGCTGTGCCCATCCTGGTAGTCATCATCCCTATCACGCTTTCGAGTGTCTGGTGGCGGATAT CCCC

### List of strains used

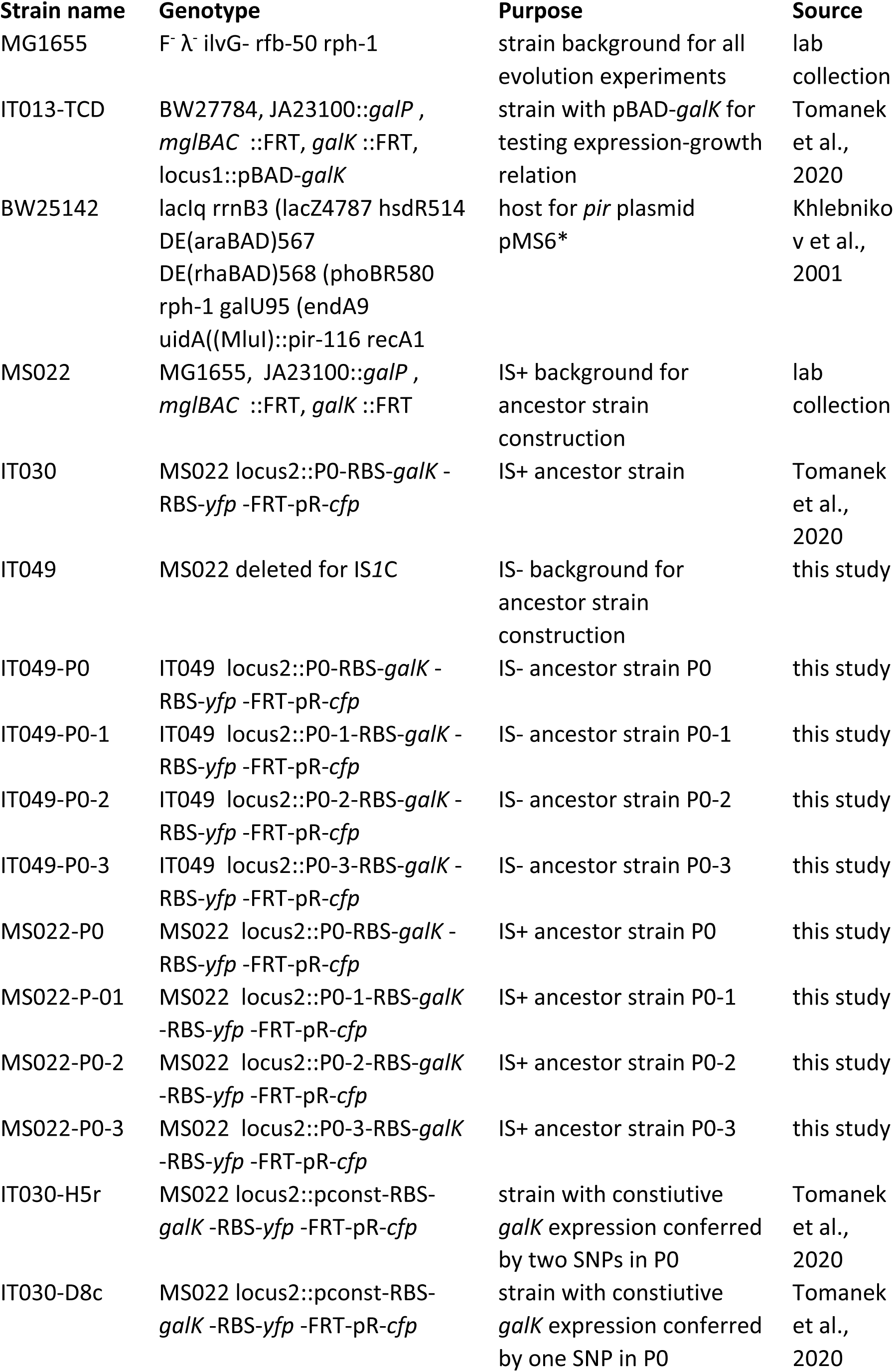

### List of primers used

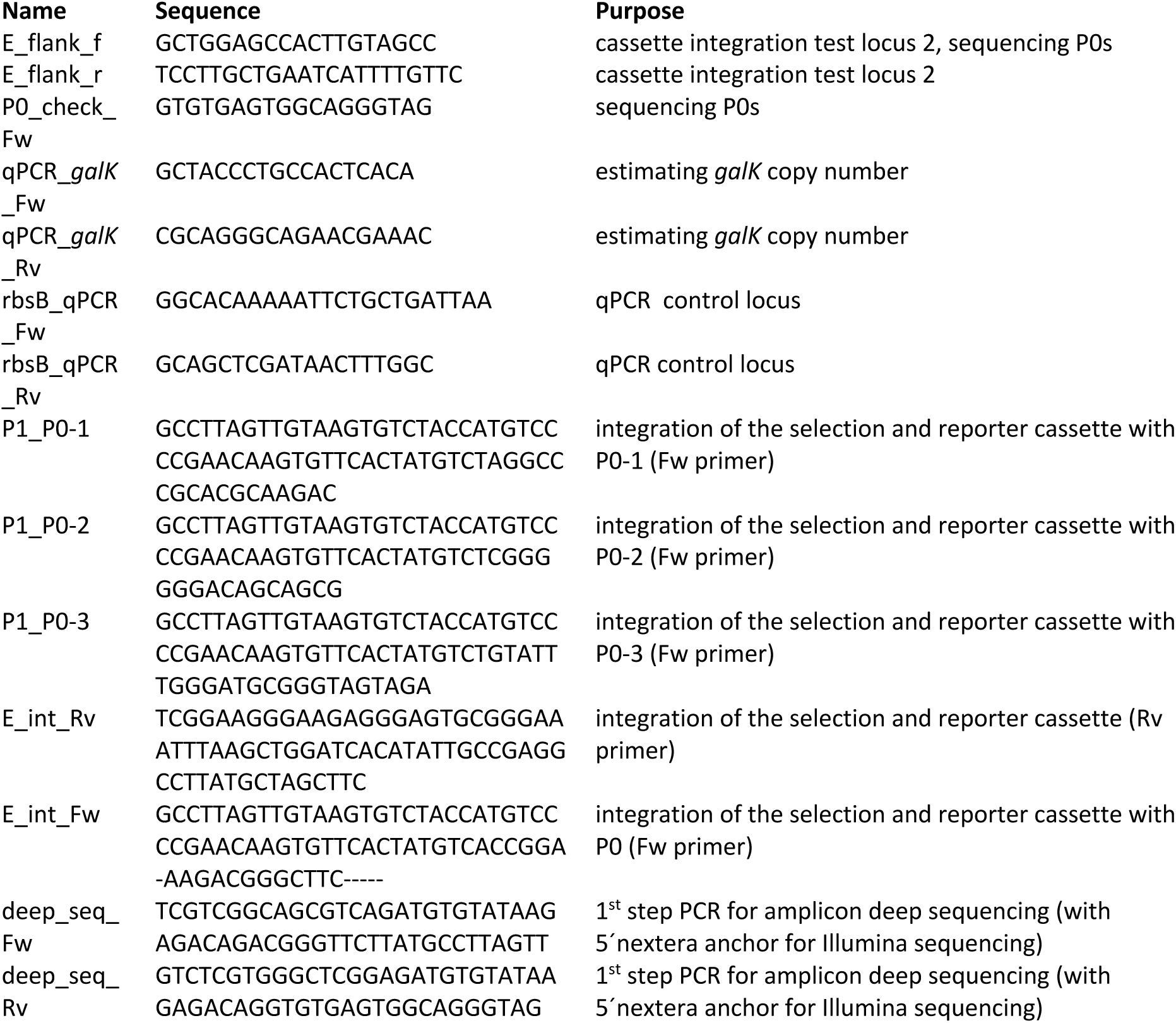

### Evolution experiments

Evolution experiments were inoculated with ancestral colonies of IS+ and IS-strains grown in 3 ml of LB medium over night, after two washing steps in M9 buffer and a dilution of 1:200.

All evolution experiments were conducted in M9 medium supplemented with 2 mM MgSO_4_, 0.1 mM CaCl_2_, 0.1% casaminoacids and carbon source at the indicated concentration (Sigma-Aldrich, St. Louis, Missouri). Bacterial cultures were grown in 200 µl liquid medium in 96-well plates and shaken in a Titramax plateshaker at 750 rpm (Heidolph, Schwabach, Germany), allowing for a total population size of ∼ 10^8^ colony forming units for the ancestral strain. Every day, populations were transferred to fresh plates using a VP408 pin replicator (V&P SCIENTIFIC, INC., San Diego, California) resulting in a dilution of ∼ 1:820 (Steinrueck and Guet, 2017), corresponding to ∼10 generations. Immediately after the transfer, growth and fluorescence measurements were performed in the overnight plates using a Biotek H1 plate reader (Biotek, Vinooski, Vermont). Thus, population phenotypes were measured every 10 generations.

### Flow cytometry experiments

Frozen evolved populations (-80°C, 15% glycerol) from day 4, day 8 or day 12 (as indicated in the figures) were pinned (1:820) into M9 buffer and put on ice until the measurement. Fluorescence was measured using a BD FACSCanto™ II system (BD Biosciences, San Jose, CA) equipped with FACSDiva software. CFP fluorescence was collected with a 450/50-nm bandpass filter by exciting with a 405-nm laser. YFP fluorescence was collected with a 510/50 band-pass filter by exciting with a 488nm laser. The bacterial population was gated on the FSC and SSC signal resulting in approximately _600_0 events analyzed per sample, out of 10,000 recorded events

### Quantitative real-time PCR

For qPCR, gDNA was isolated from overnight cultures grown in the respective evolution medium inoculated by single evolved colonies using Wizard Genomic DNA purification kit (Promega, Madison, Wisconsin). We performed qPCR using Promega qPCR 2x Mastermix (Promega, Madison, Wisconsin) and a C1000 instrument (Bio-Rad, Hercules, California). To quantify the copy number of samples of an evolving population, we designed one primer pair within *galK* (target) and one primer within *rbsB* as a reference, which lies outside the amplified region. We compared the ratios of the target and the reference loci to the ratio of the same two loci in the single copy control. Using dilution series of one of the gDNA extracts as template, we calculated the efficiency of primer pairs and quantified the copy number of *galK* in each sample employing the Pfaffl method, which takes amplification efficiency into account (Pfaffl, 2001). qPCR was performed in three technical replicates.

### Measurement of colony fluorescence

Evolving populations were pinned onto LB agar supplemented with 1% charcoal and imaged using the macroscope set up (https://openwetware.org/wiki/Macroscope) (Chait *et al*., 2010). To obtain median colony YFP and CFP fluorescence intensity, a region of interest was determined using the ImageJ plugin ‘Analyze Particles’ (settings: 200px-infinity, 0.5-1.0 roundness) to identify colonies on 16-bit images with threshold adjusted according to the default value. The region of interest including all colonies was then used to measure intensity.

### Amplicon deep sequencing of P0

Frozen samples of evolved populations were diluted 1:10 into 100 µl of LB and grown for 5 hours (37°C, shaking) to increase cell numbers prior to DNA extractions. Columns 1-4 (populations A1, B1, C1… F4, G4, H4), 5-8 (populations A5, B5, C5… F8, G8, H8) and 9-12 (populations A9, B9, C9… F12, G12, H12) of each 96 well plate were pooled prior to DNA extraction using Wizard Genomic DNA purification kit (Promega, Madison, Wisconsin). The P0 region including the beginning of *galK* was amplified for 25 PCR cycles using primers deep_seq_Fw and deep_seq_Rv carrying 5’ adaptors for Illumina sequencing. In parallel, PCR reactions were performed for 35 cycles to confirm bands on a gel. Illumina sequencing was carried out by Microsynth (Balgach, Switzerland). We note that our amplicon libraries of P0 were contaminated with reads carrying the sequence of P0-2, which we had prepared for sequencing in parallel (Supplementary Fig. 4). We therefore excluded all reads of P0-2 for our analysis of P0 and do not report the result of the P0-2-specific samples as they could not be trusted.

Reads of P0 were analyzed using a custom R script. Briefly, we defined four sequence motifs of each 39 bp length, which represented the ancestral P0 sequence and the same region with known adaptive SNPs included (T>A, C>T or both). We counted the number of reads with ancestral or evolved 39bp motif in all samples, including those of control populations evolved in the absence of galactose. We also counted the number of reads with an ancestral *galK* sequence motifs spanning 39bp as well as the mean number of reads that carry a 39-bp ancestral *galK* sequence motif with one single SNP.

## Author contribution

C.C.G. and I.T. designed study, I.T. carried out experiments and analyzed data, C.C.G. and I.T. wrote the manuscript.

---

**Supplementary Table 1.** Sequencing and phenotypic analysis of all YFP+ IS+ populations evolved in 0.01% galactose (Fig. 4D - red triangles). Increase in fluorescence relative to ancestral (anc) phenotype indicated by YFP+ and CFP+. Results shown for day 12 populations unless otherwise noted (d4, d8).

| Population | seq (all YFP+) | flow cytometry phenotype | agar streak | comment |
| --- | --- | --- | --- | --- |
| A6 | -30T>A | YFP+, v. few CFP+ (mixed populations) | YFP+, few CFP+ |  |
| B1 | -30T>A, -37C>T ("mutation H5") | YFP+, CFP+ (mixed populations) | few YFP+, few CFP+, mixed pop |  |
| B2 | -30T>A | YFP+ | YFP+, v. few CFP+ |  |
| C1 | -30T>A | YFP+ (d12) | YFP+, v. few CFP+ |  |
| C9 | - | ancestral YFP (d8), only CFP+(d12) | - | incorrectly classified as YFP+ (Fig. 4D – grey triangle) |
| D2 | -30T>A | YFP+ (d12) | YFP+ only |  |
| D9 | anc | YFP+ (d8, d12) | YFP+ only |  |
| E9 | anc | anc | - |  |
| E10 | -30T>A | YFP+ (d12) | YFP+ only |  |
| F6 | - | YFP+ (d4), CFP+(d12) | - | YFP+ at d8, then amplified population (Fig. 5D - blue triangle) |
| F10 | -30T>A | YFP+, CFP+, anc (mixed populations) | YFP+,CFP+, mixed pop | qPCR confirmed |
| G1 | -30T>A | YFP+(d4-8), v. few CFP+ (d12) | YFP+, v. few CFP+ |  |
| G12 | -30T>A | YFP+ (d8), CFP+ (d12) | YFP+, only | FACS CFP+ carry-over |

**Supplementary Table 2.** Mutations of P0-2 underlying increased YFP fluorescence in IS+ and IS-populations evolved in 0.1% galactose.

| IS + clones |  | IS – clones |  |
| --- | --- | --- | --- |
| **P02-A11** | -131_-144del | **P02-A7** | -100C>T |
| **P02-B10** | -122_-134del | **P02-H12** | -100C>T |
| **P02-F4** | -100C>T | **P02-C3** | -100C>T |
| **P02-F4** | -100C>T, poor quality read | **P02-H9** | -122_-134del |
|  |  | **P02-F2** | -122_-134del |
|  |  | **P02-D1** | -100C>T |
|  |  | **P02-E2** | -100C>T |
|  |  | **P02-A1** | bigger band, maps to *insD1* coding sequence |
|  |  | **P02-E5** | -41del |
|  |  | **P02-C5** | 201bp deletion leaving 20bp of P02 |
|  |  | **P02-H5** | 201bp deletion leaving 20bp of P02 (7 different kinds of mutations) |

**Supplementary Figure 1.**
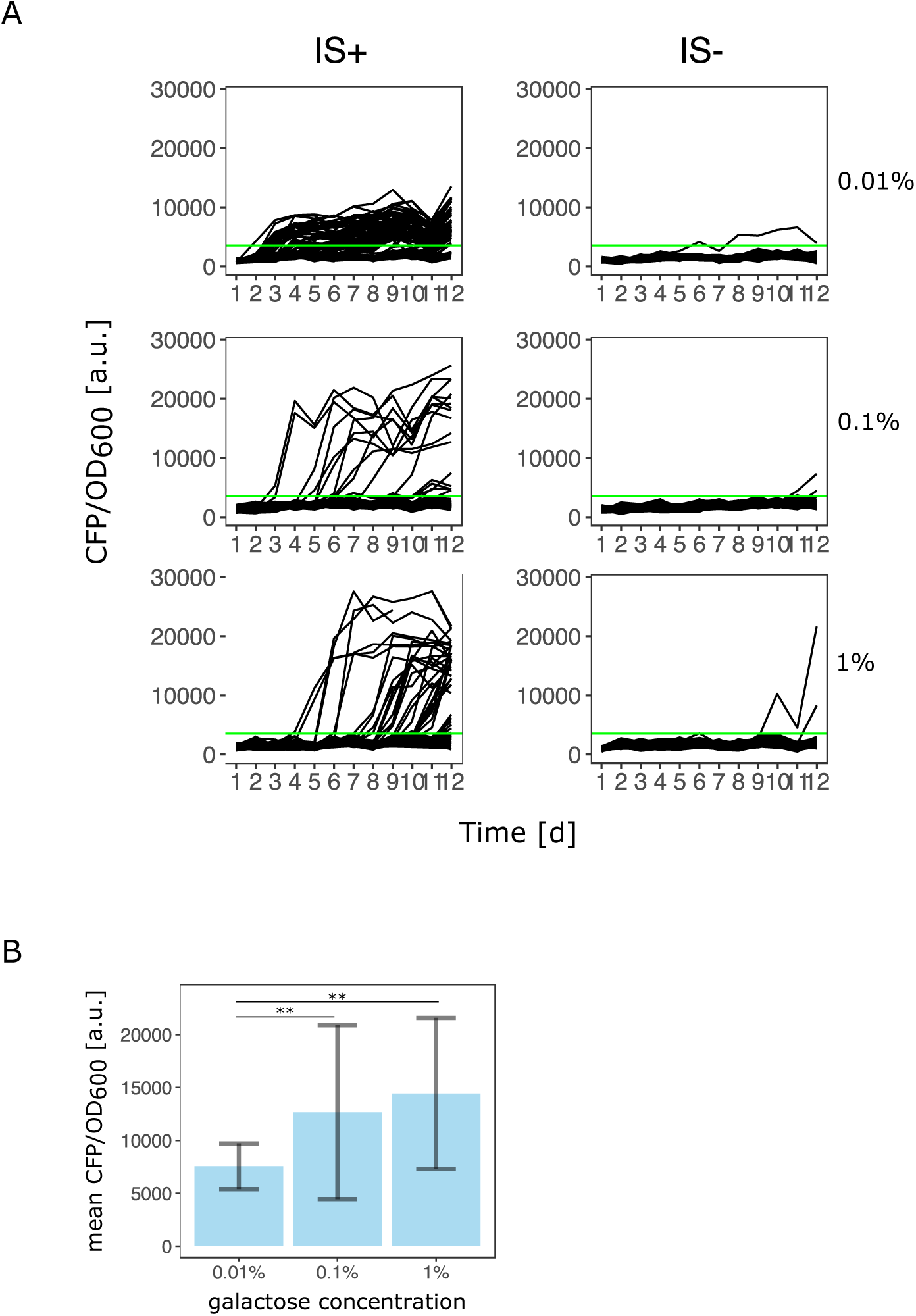
Number of amplified populations and their copy-number depends on the gene expression demand of the environment. **A**. Data replotted from Fig. 2A. Green line indicates threshold to classify as population as amplified (CFP/OD_600_ exceeds the mean ancestral CFP/OD_600_ by four standard deviations). **B**. Using the same threshold, mean CFP/OD_600_ fluorescence as a proxy for copy-number of all evolved populations is shown for 0.01%, 0.1% and 1% galactose (68, 19 and 34 populations for low, intermediate and high galactose, respectively). p-values (two-sided *t*-test): 3.6*10^−6^ (between 0.01% and 1% gal) and 3.10^−2^ (between 0.01% and 0.1% galactose).

**Supplementary Figure 2.**
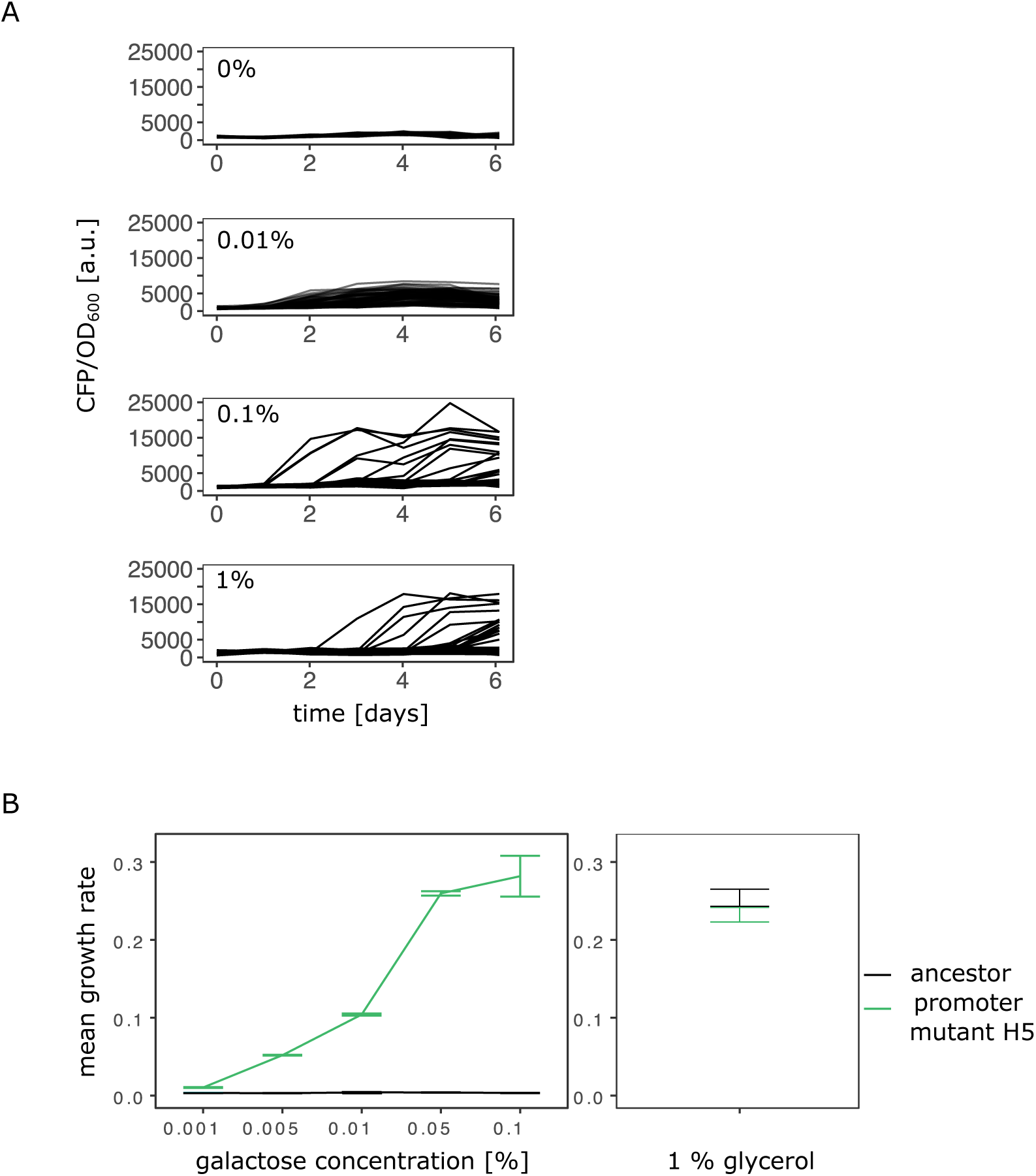
Evolutionary dynamics depend on galactose concentration. **A**. Additional evolution experiment with daily measurements of normalized CFP fluorescence as a proxy for gene copy-number of 96 populations of the IS+ strain growing in three different galactose concentrations (% indicated next to the plots), as well as in the absence of galactose (control). **B**. Growth rate in minimal medium with increasing concentrations of galactose (left panel) as well as glycerol (control, right panel) of strain H5 with two SNPs in P0 (-30T>A and -37C>T) and the ancestral strain. Error bars represent the standard deviation of four (galactose) and five (glycerol) replicates, respectively.

**Supplementary Figure 3.**
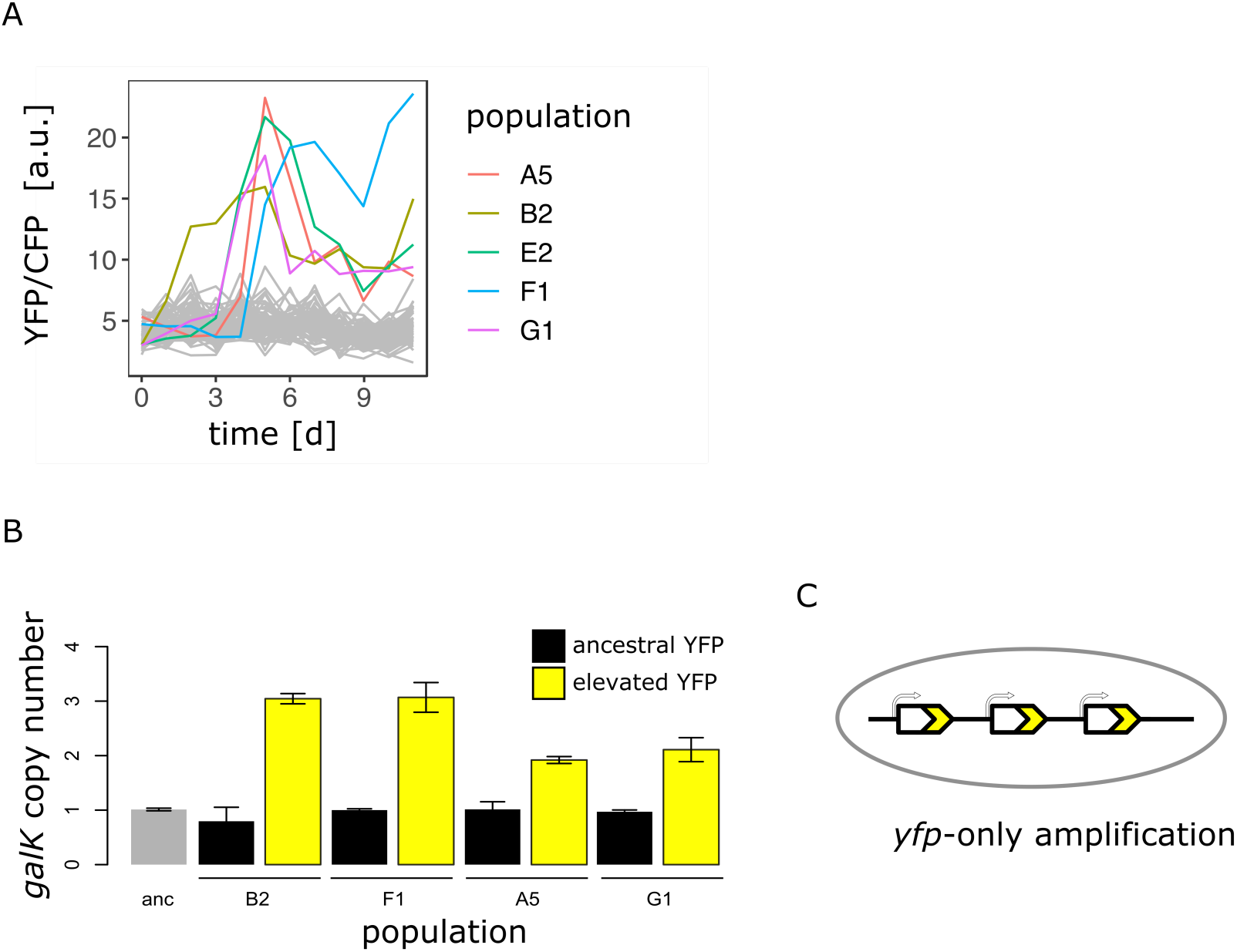
YFP-only amplifications occur in IS-populations evolved in 0.1% galactose. **A**. Normalized YFP fluorescence as a proxy for *galK* expression of 96 populations in the IS-strain growing in 0.1% galactose. Populations with increased YFP fluorescence are highlighted. ***B***. *GalK* copy-number of the YFP+ IS-populations evolved in 0.1% galactose shown in (**A**) as estimated by qPCR. For each population, genomic DNA of one colony with ancestral (black bars) and one with increased YFP (yellow bars) fluorescence was analyzed. **C**. Scheme of *galk-yfp*-only amplification with a duplication junction upstream of the *cfp* gene.

**Supplementary Figure 4.**
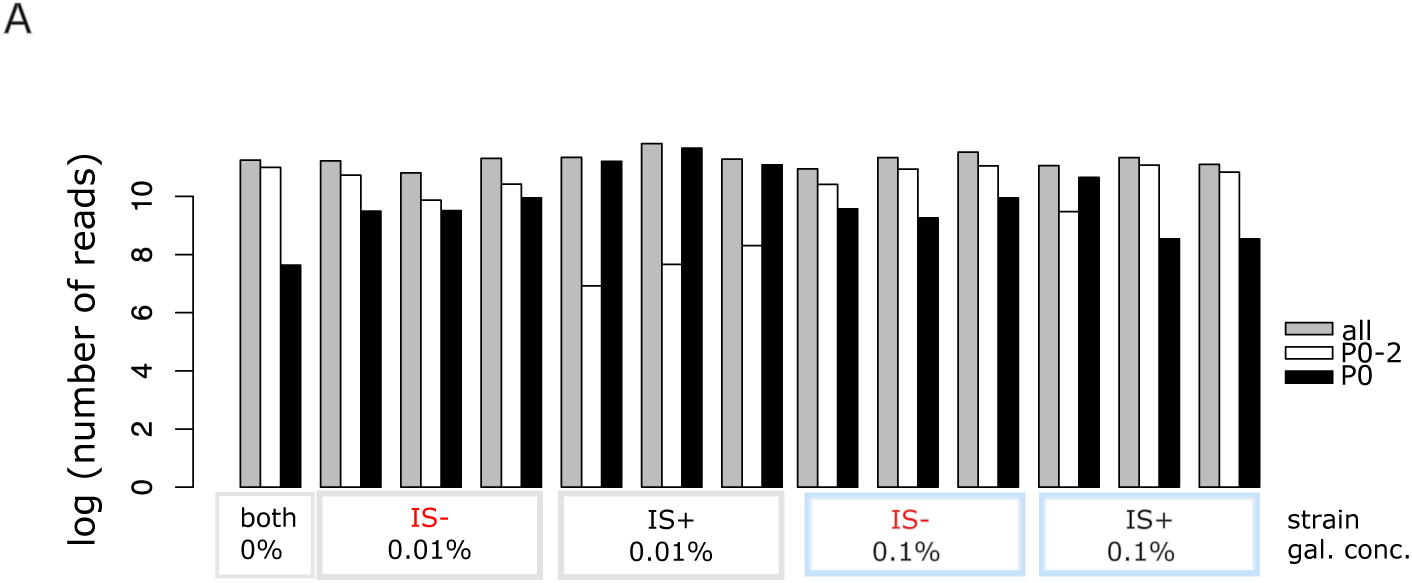
Total number of sequencing reads for all replicates. **A**. Log plot of total read numbers showing contamination of P0 amplicons with P02 amplicons stemming from pooled samples of the 0.1% galactose populations of both promoter sequences (blue rectangles; see Methods).

**Supplementary Figure 5.**
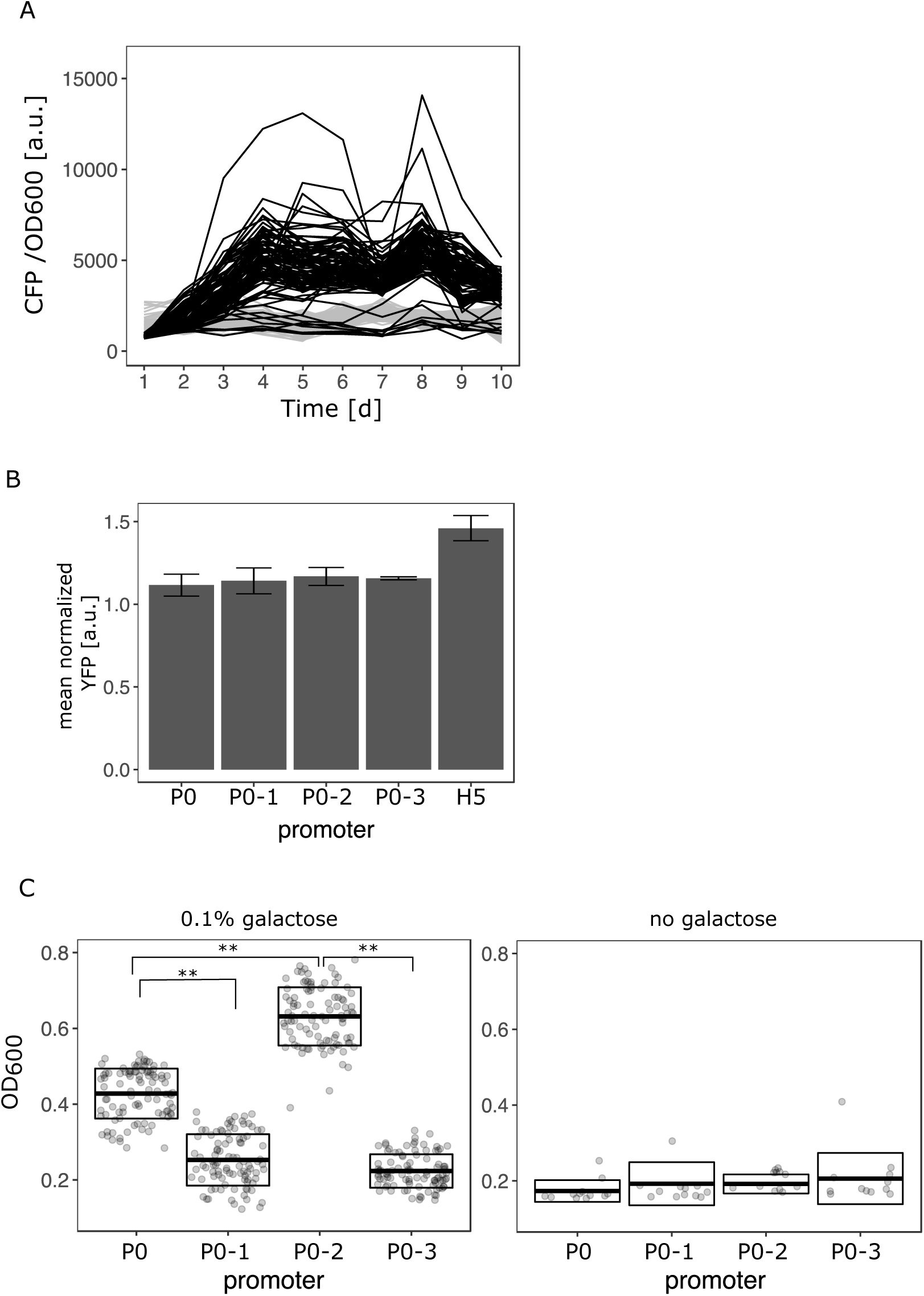
Rapid amplification of IS+ populations with P02. **A**. CFP/OD_600_ as a proxy for copy-number plotted over the course of the evolution experiment for IS+ with P0-2 populations in 0.1% galactose and control populations in 0% galactose (grey). **B**. Flow cytometry measurement of YFP fluorescence intensity as a proxy for *galK* expression of IS-strains harboring the four random promoter sequences as well as a P0 with adaptive SNPs as a comparison (“H5”; indicated at the bottom of the figure), respectively, normalized to a strain without fluorescence marker. Error bars represent the standard deviation of three biological replicates. **C**. End-point OD_600_ (‘yield’) of IS-populations carrying P0, P0-1, P0-2 and P0-3 after 24h of growth in 0.1% galactose (left panel) and in the absence of galactose (right panel). Boxes indicate the mean and standard deviation of 96 populations (left panel) and 12 populations (right panel), respectively. Asterisks indicate a significant difference between mean OD_600_ (two-sided *t*-test, p <0.0001).

